# Review of methods to derive the heartbeat-evoked potential: past practices and future directions

**DOI:** 10.1101/2024.07.23.604405

**Authors:** Rania-Iman Virjee, Rohan Kandasamy, Sarah N Garfinkel, David W Carmichael, Mahinda Yogarajah

## Abstract

The heartbeat-evoked potential (HEP) is an implicit, electrophysiological marker of cortical heartbeat processing and interoception, with increasing clinical relevance. However, on the scalp, HEP are low-amplitude signals mixed with cardiac field artefacts (CFA), requiring signal processing pipelines to separate HEP from CFA. This review evaluates current analytical approaches, addresses methodological gaps in HEP pipelines, and examines the impact of key parameter choices.

HEP processing methods/parameters used in EEG (N=101) and MEG (N=10) studies were investigated, focusing on the effects of HEP window, electrodes, filters, independent component analysis (ICA) and artefact subspace reconstruction (ASR), on HEP extraction, using Temple University’s normal scalp EEG data.

EEG and MEG studies revealed clear inconsistencies in HEP parameter use and reporting. ASR-20 (ASR threshold for artifact identification) performed comparably to ICA for artefact removal, supporting its potential real-time EEG applicability. Epoch rejection, a HEP quality metric, appeared equivalent between ICA and ASR-20 after artefact removal. Linear Mixed Model analysis identified significant effects of RR interval, maximum epoch amplitude, HEP window and baseline correction start time on measured HEP amplitude.

Publications should report critical values for reliable HEP extraction, emphasising the need for standardised methods to enhance study comparability and reproducibility.

## 1. Introduction

Interoception is the process by which the nervous system senses, interprets, and integrates signals originating from within the body (e.g. hunger, thirst, cardiac and respiratory signals), providing a moment-by-moment mapping of the body’s internal landscape across conscious and unconscious levels (Khalsa et al., 2018). This is essential for homeostasis and allostasis. Indeed, the heart continuously and cyclically communicates with the brain, impacting cognitive and emotional processes (Park and Blanke, 2019). First investigated by Schandry et al. (1986) and Jones et al. (1986), the heartbeat-evoked potential (HEP) is a cortical brain response that is time-locked to each heartbeat, which has been proposed as an implicit, electrophysiological marker reflecting the cortical processing of heartbeats (Kumral et al., 2022), and more broadly interoception. Since these initial publications in the early 1980s, HEP has gradually gained more attention in the literature, with publications rising to approximately 90 in the year 2021 (Shamsaei et al., 2010).

HEP appears to be a meaningful clinical measure (Suksasilp and Garfinkel, 2022). For example, HEP has been shown to have trait variation between groups such as in the ageing population (Kamp et al., 2021), and in groups with psychiatric conditions characterised by emotion dysregulation, including borderline personality disorder (BPD) (Flasbeck et al., 2020), and generalised anxiety disorder (GAD) (Pang et al., 2019; Schmitz et al., 2021). Other studies have shown state differences in HEP, including relationships with interoceptive processing (Baranauskas, Grabauskaitė and Griškova-Bulanova, 2017; Desmedt et al., 2023; Hodossy et al., 2021), as well as in the periods preceding functional seizures (FS) (Elkommos et al., 2023). Other studies aiming to validate brain stimulation techniques to interfere with interoceptive processes (Pollatos et al., 2016) and novel behavioural tasks to assess interoceptive processing in infants (Maister et al., 2017) have also utilised HEP, although these have been debated by the community (Coll, Penton, & Hobson, 2017).

Despite these promising earlier indicators of HEP utility, a fundamental inherent constraint is its susceptibility to contamination from direct electrical potential and associated magnetic flux when measuring it with electroencephalogram (EEG) or magnetoencephalography (MEG). This contamination arises from the electrophysiological activity of the heart which generates an electric field (i.e. the cardiac field artefact - CFA) that spreads throughout the whole body including the scalp (Arnau et al., 2023). Consequently, removing the CFA from HEP presents a significant challenge. This challenge stems from the limited capability to precisely measure the cardiac field and the incomplete understanding of its propagation towards the scalp (Yuan et al., 2007).

Studies have suggested various physiological sources underlying HEP (Park and Blanke, 2019). Previous work suggests that the physiological pathways are multifaceted and complex. HEP components can be noted in multiple cortical regions and temporal intervals, which may relate to distinct afferent signals from the carotid arteries and baroreceptors at the aortic arch (Gray et al., 2007; Garfinkel and Critchley, 2016), the cardiac afferent neurons (Tahsili-Fahadan and Geocadin, 2017), somatosensory pathways (Khalsa et al., 2009; Kern et al., 2013) and cortical neuro-vascular coupling (Kim et al., 2016). To date, studies have reported HEP in numerous sensor locations: frontal (Canales-Johnson et al., 2015; Babo-Rebelo et al., 2016a), central (Montoya et al., 1993; Petzschner et al., 2019) and parietal channels (Pollatos and Schandry 2004; Babo-Rebelo et al., 2016a; Kern et al. 2013). This variability in HEP topography is thought to be generated by various methodological considerations and the widespread distribution of interoceptive processing in the brain including the insula, cingulate cortex, somatosensory cortex and amygdala. Variation in experimental design, such as the positioning of the reference, the number of electrodes, and the performance of the EEG amplifier can all in turn affect HEP topographies (Park and Blanke, 2019). The absence of control conditions in some experimental designs also contributes to the complexity in understanding physiological pathways underlying HEP (Coll, Penton and Hobson, 2017). As Coll et al. (2021) identified, a systematic meta-analysis of HEP research is essential to render the use of HEP as a reliable, powerful and well-grounded indicator of cortical interoceptive processing of cardiac signals and to establish consistent processing methodologies to use HEP in clinical and research settings (Coll et al., 2021; Suksasilp and Garfinkel, 2022).

The aims of the review are threefold. Firstly, to better understand HEP processing methods currently used in both EEG and MEG studies. This review aims to explore and quantify variability in the methodology and parameters used to record the heartbeat-evoked potential. Secondly, based on the HEP processing pipelines found in the literature, a sensitivity analysis will be performed to demonstrate the potential impact of different pipeline parameters on HEP values using EEG. Finally, this review aims to take an initial step towards fostering consensus about optimal and standardised HEP processing methods through an informed assessment of different methods to date and their impact on HEP signal.

## 2. Materials and methods

### 2.1. Scoping review search strategy

The literature search involved two electronic databases, Web of Science and PubMed, which yielded 1001 potential eligible studies. This was reduced to 124 studies (EEG and MEG studies) after removal of duplicates and exclusion of studies (figure 1). This search spanned from 1993 to 2023. A series of different combinations of key words were used for the literature search including Boolean operators (‘AND’ and ‘OR’) (table 1). The two studies using intracranial EEG (iEEG) were excluded as they did not provide enough data to support robust conclusions.

**Figure 1.**
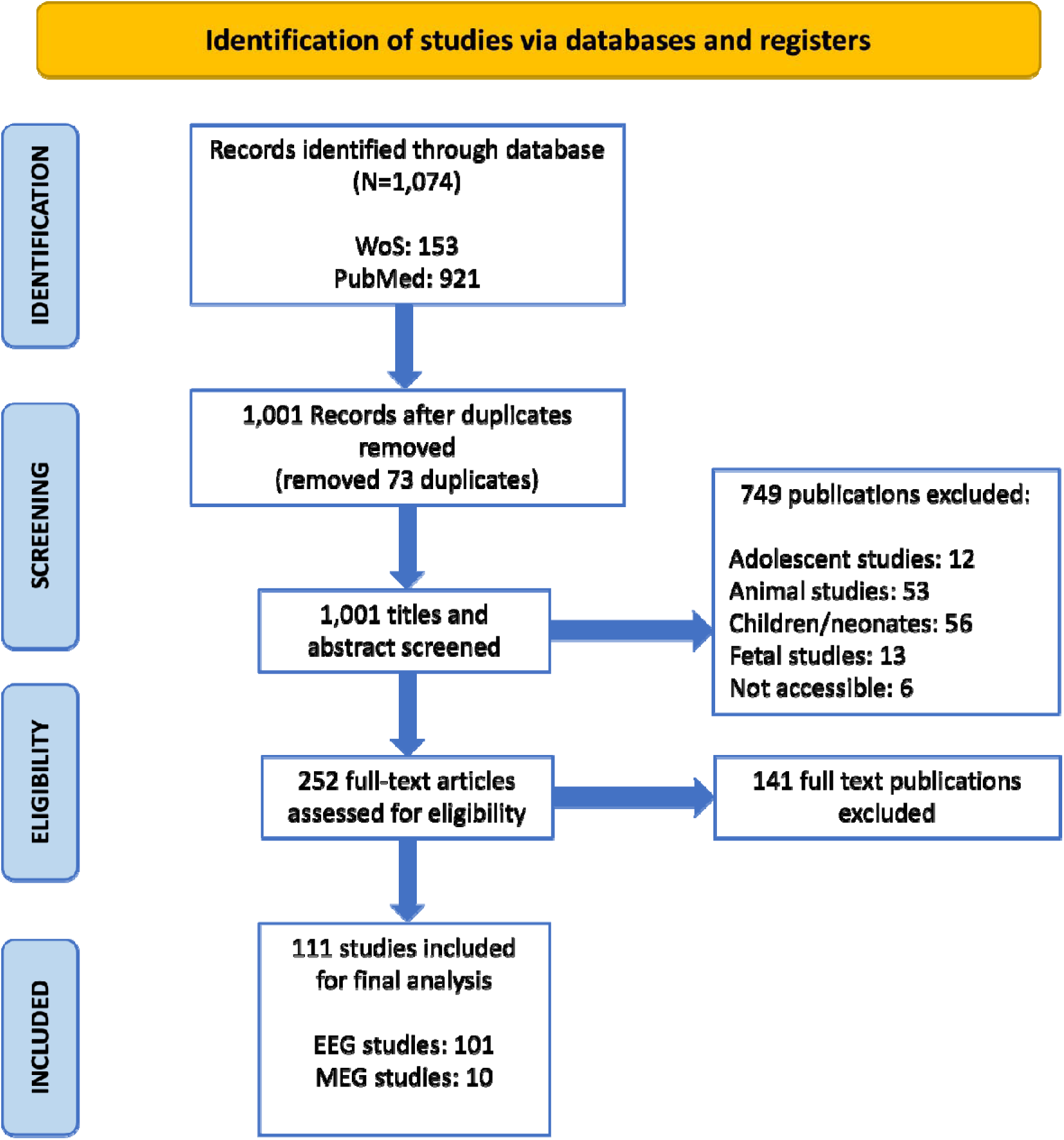
PRISMA flow diagram demonstrating the total number of publications included and removed at the different stages and the reasons for exclusion.

**Table 1.** Search strategies used in different databases.

| PubMed | Or | And |
| --- | --- | --- |
|  | "heart" or "heart rate" | "evoked potentials" |
|  | "heart" | "evoked potential" |
|  | "HEP" | "electroencephalography" or<br>"electroencephalogram" |
|  | "EEG"<br>or "magnetoencephalography"<br>or "magnetoencephalogram" |  |
|  | "MEG" |  |
| Web of Science | "heartbeat evoked" |  |

### 2.2. Inclusion and exclusion criteria

Studies were included if they used adult human HEP derived from EEG or MEG. Exclusion criteria encompassed animal studies (rabbits, dogs, cats, rats), children and neonatal studies, adolescent studies, foetal studies, and other studies not related to HEP (such as other event-related potentials, epilepsy studies, viruses, or protein) or inaccessible studies.

### 2.3. Data collection and analysis

Full-text screening and data extraction was performed by the author and an independent reviewer (co-author: R.K.) flagged any inconsistencies during data extraction. Current processing techniques and parameters reported in the literature were examined and are listed below. The data extracted from the studies underwent collation, synthesis, and analysis based on the following variables and their definitions (table 2, figure 2):

A. Low-pass and high-pass filter cut-off frequencies in Hertz (Hz)
B. Epoch timeframes in seconds (s)
C. Epoch relative to R wave or T wave
D. Epoch amplitude rejection in microvolts (µV)
E. HEP window in milliseconds (ms)
F. Baseline correction time frame in milliseconds (ms)
G. Minimum RR interval in milliseconds (ms)
H. CFA removal methodology
I. Electrodes used for EEG analysis

**Table 2.**
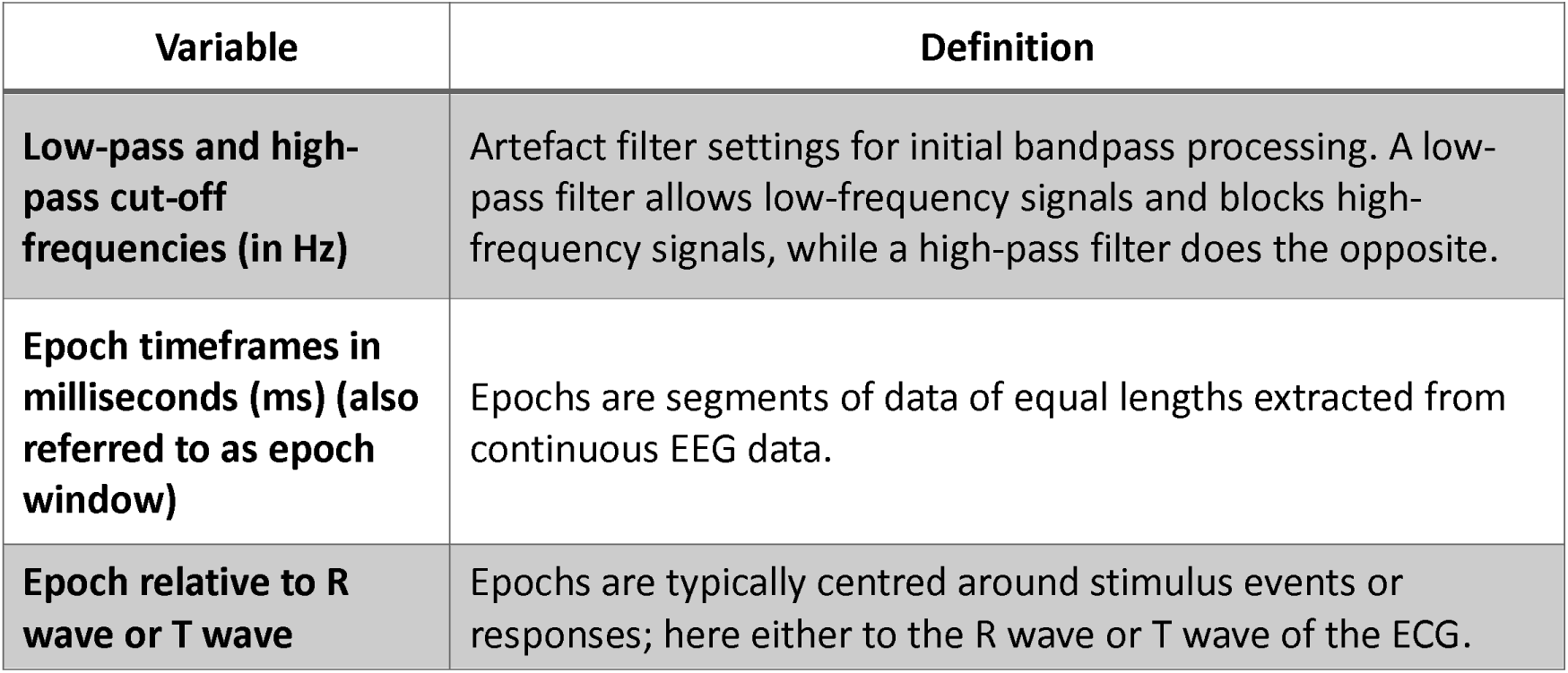
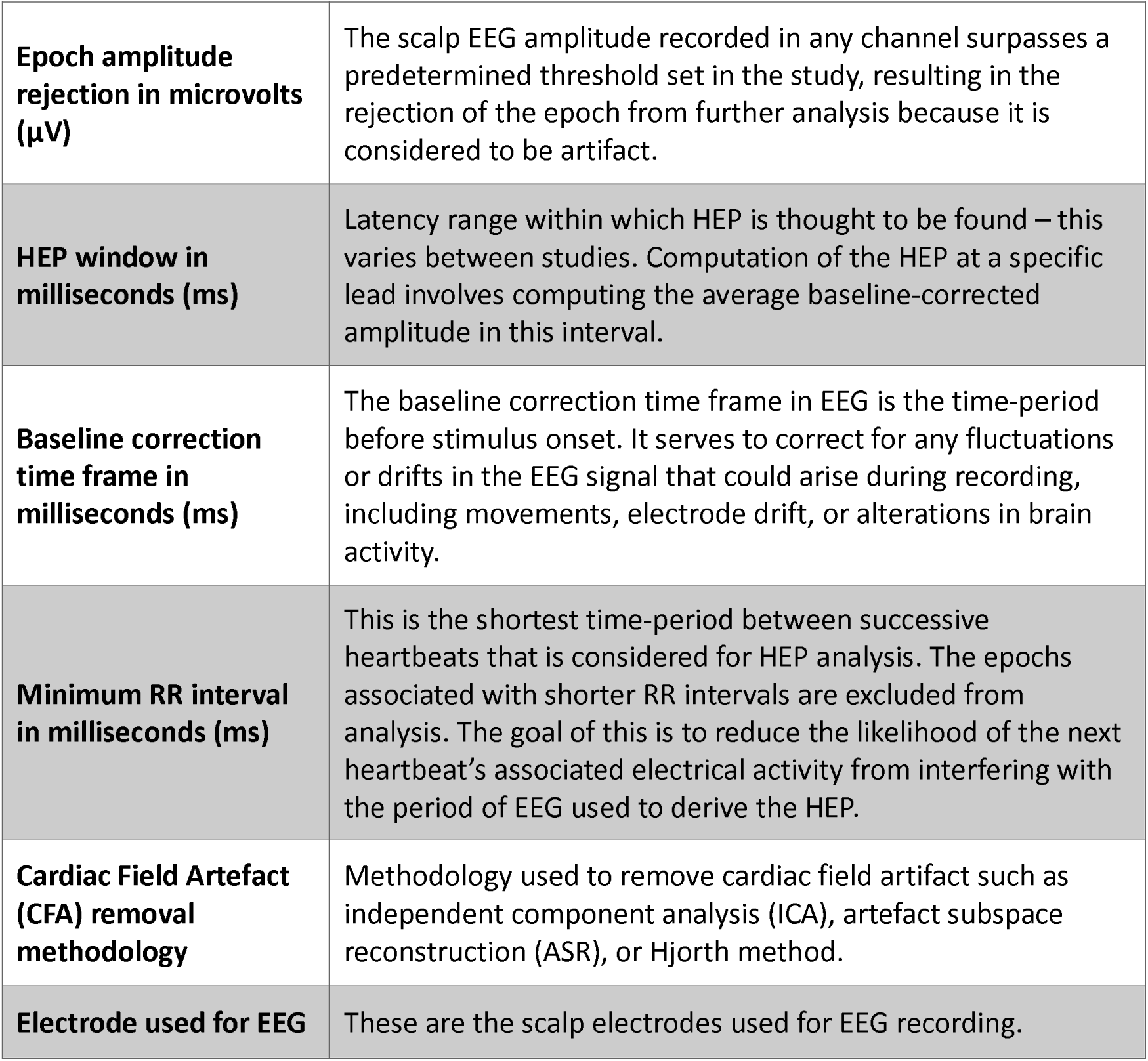
Variables extracted from the literature and their definitions. Independent component analysis (ICA) is a signal processing technique which separates EEG signal into independent source components. Some of these components may correspond to artefacts such as muscle activity or eye movements. Artefact components are then identified, removed manually and computationally. Artefact subspace reconstruction (ASR) requires an initial calibration using segments (detected manually or automatically) of relatively clean baseline EEG data. ASR is designed to handle non-stationary/non-repeating artefacts using a sliding window and principal component analysis to identify and remove high-amplitude signal components. The three Hjorth parameters – activity (variance of the signal), mobility (mean frequency of the power spectrum), and complexity (estimation of the bandwidth of the signal) - are calculated using time-domain signal processing approaches, which offer a statistical analysis of the signal’s attributes. These parameters are used to identify and remove artefacts.

| Variable | Definition |
| --- | --- |
| Low-pass and high-pass cut-off frequencies (in Hz) | Artefact filter settings for initial bandpass processing. A low-pass filter allows low-frequency signals and blocks high-frequency signals, while a high-pass filter does the opposite. |
| Epoch timeframes in milliseconds (ms) (also referred to as epoch window) | Epochs are segments of data of equal lengths extracted from continuous EEG data. |
| Epoch relative to R wave or T wave | Epochs are typically centred around stimulus events or responses; here either to the R wave or T wave of the ECG. |
| **Epoch amplitude rejection in microvolts ($\mu\text{V}$)** | The scalp EEG amplitude recorded in any channel surpasses a predetermined threshold set in the study, resulting in the rejection of the epoch from further analysis because it is considered to be artifact. |
| **HEP window in milliseconds (ms)** | Latency range within which HEP is thought to be found – this varies between studies. Computation of the HEP at a specific lead involves computing the average baseline-corrected amplitude in this interval. |
| **Baseline correction time frame in milliseconds (ms)** | The baseline correction time frame in EEG is the time-period before stimulus onset. It serves to correct for any fluctuations or drifts in the EEG signal that could arise during recording, including movements, electrode drift, or alterations in brain activity. |
| **Minimum RR interval in milliseconds (ms)** | This is the shortest time-period between successive heartbeats that is considered for HEP analysis. The epochs associated with shorter RR intervals are excluded from analysis. The goal of this is to reduce the likelihood of the next heartbeat's associated electrical activity from interfering with the period of EEG used to derive the HEP. |
| **Cardiac Field Artefact (CFA) removal methodology** | Methodology used to remove cardiac field artifact such as independent component analysis (ICA), artefact subspace reconstruction (ASR), or Hjorth method. |
| **Electrode used for EEG** | These are the scalp electrodes used for EEG recording. |

**Figure 2.**
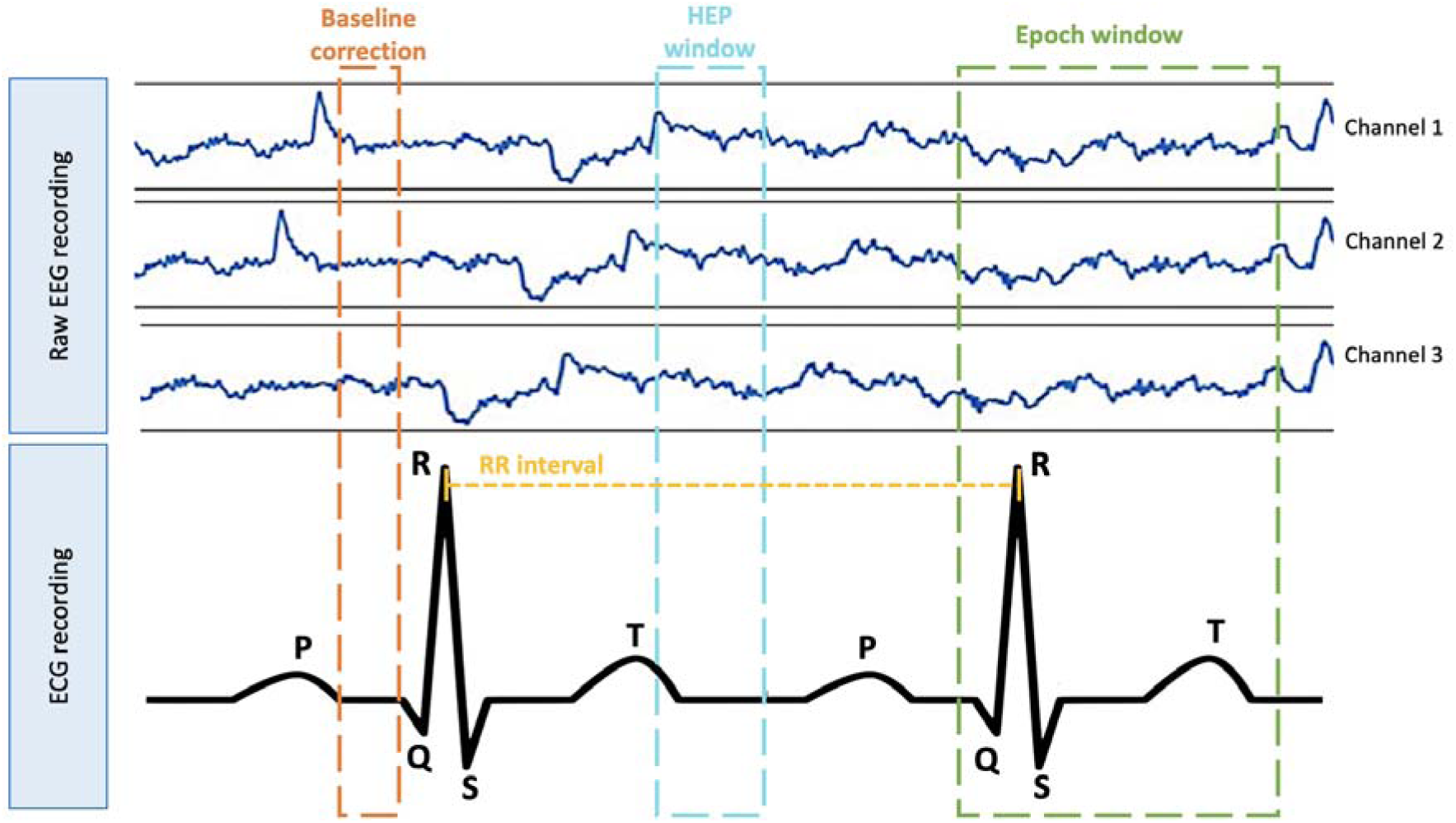
Diagram representing key elements of the EEG and ECG signals to process EEG and derive HEP. Note that the placement of the HEP window, epoch window and baseline correction window are prototypical and may vary in position relative to components on the ECG trace, depending on the study’s methodology.

As only two studies used the T wave for epoch centring, this parameter was not further investigated. Following data extraction, statistical analysis was performed using Python version 3.9 and the MNE python package version 1.7.0 (Larson, 2024).

### 2.4. Processing pipeline methodology

The second part of this review aims to contribute to building consensus about optimal and standardised HEP processing methods through an informed assessment of different methods used to date in the literature and their impact on HEP signal. This was achieved by developing a signal processing pipeline for HEP. Different processing methods extracted from the literature were implemented and tested on 97 randomly selected available scalp EEG recordings (normal EEG studies from the Temple University Hospital EEG (TUH EEG) Corpus (Obeid and Picone, 2016)) to explore how different methods and parameters affect the HEP extraction process and results. These were clinically sourced EEG recordings, using the 10-20 montage, and reported by experts. Factors explored for their effect on HEP included the same variables identified during data extraction (section 2.3). To do this, relatively clean sections of EEG were clipped by hand (using EDFbrowser software version 1.4), then pre-processed with variable applications of artefact-subspace-reconstruction (ASR) or independent-component-analysis (ICA). Epochs were generated with various values of minimum RR threshold, maximum epoch amplitude, and filter settings (figure 3). HEP was then computed for each set of generated epochs. The variability of the HEP between epochs for a given set of parameters was analysed following the values described in Table 3 and in Figure 3. This was achieved for different pre-processing methods: ICA without CFA removed, ASR at different thresholds (5, 10, and 20) and uncorrected raw EEG (figure 3). ICA did not reliably identify a component containing CFA, therefore ICA with CFA removal was not included in the analysis. This could be due to having only 20 EEG leads for analysis, and hence a maximum of 20 components, limiting the ICA algorithm’s ability to locate and remove artefacts.

**Figure 3:**
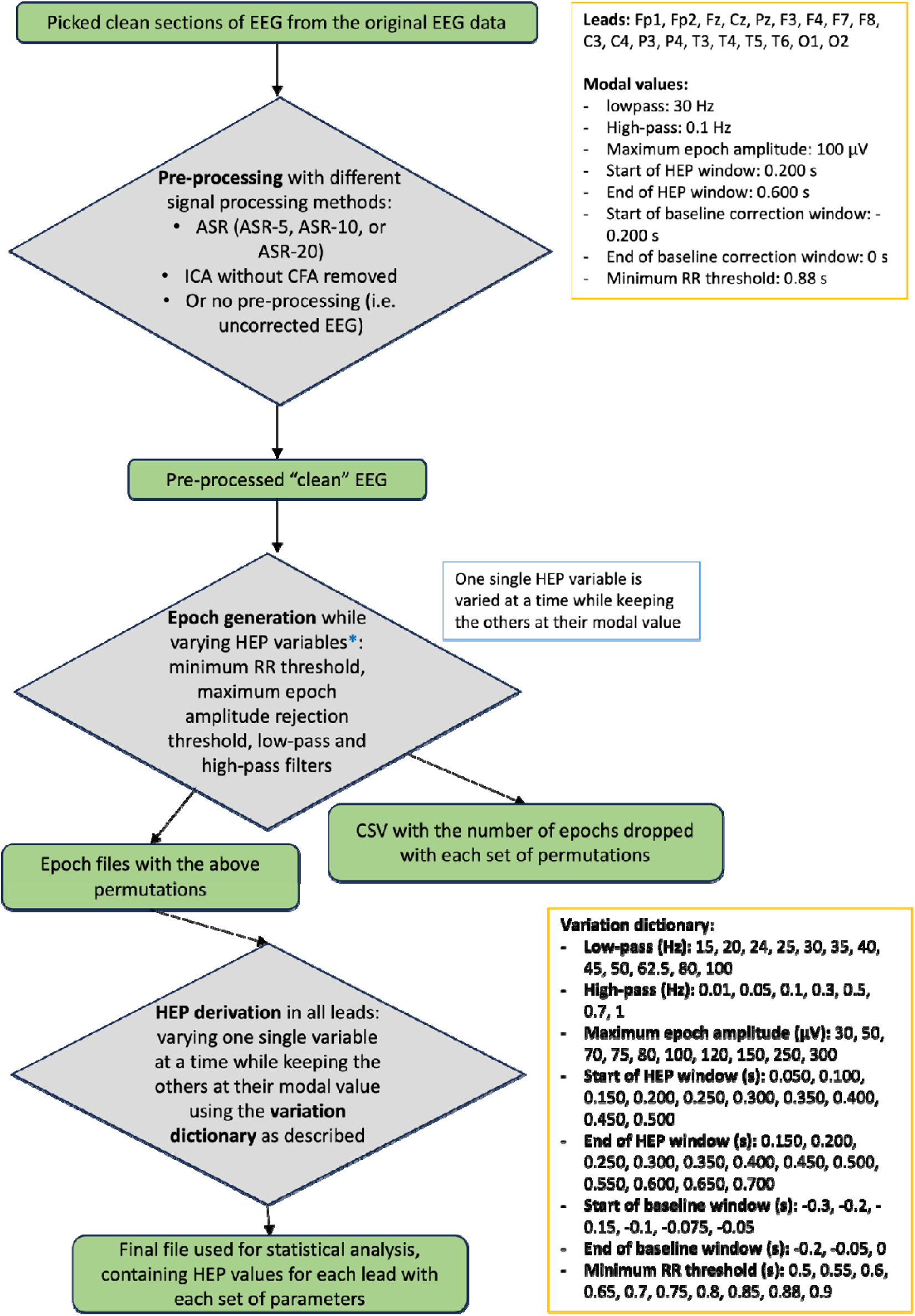
Flowchart illustrating the data processing methodology with values used for analysis and HEP computation and EEG channels considered. CSV – comma separated value files.

**Table 3.** Snapshot of values for the pre-processing variables extracted from the literature review. These were used to compute the HEP in the analysis. Values in bold were identified as the modal values in the literature (complete list of values and details in Figure 3).

| High pass (in Hz) | Low pass (in Hz) | Maximum epoch<br>amplitude (in $\mu\text{V}$) | Minimum RR<br>interval (in seconds) |
| --- | --- | --- | --- |
| 0.01 | 15 | 30 | 0.5 |
| 0.05 | 20 | 50 | 0.55 |
| **0.1** | 24 | 70 | 0.6 |
| 0.3 | 24 | 75 | 0.65 |
| 0.5 | **30** | 80 | 0.7 |
| 0.7 | 35 | **100** | 0.75 |
| 1 | 40 | 120 | 0.8 |
| - | 45 | 150 | **0.88** |
| - | 50 | 250 | 0.9 |
| - | 62.5 | 300 | - |
| - | 80 | - | - |
| - | 100 | - | - |

A sensitivity analysis was conducted to assess which variables have the greatest effect on HEP. For this, independent variables were varied (low-pass and high-pass for filtering settings, baseline start and baseline end for the baseline window, start of HEP window and end of HEP window for the HEP window, minimum RR interval and maximum epoch amplitude threshold) and the effect on dependent variables (HEP at Fp1, Fp2, Fz, Cz, Pz, F3, F4, F7, F8, C3, C4, P3, P4, T3, T4, T5, T6, O1, O2) was recorded in each filtering method (ASR 5, ASR 10, ASR 20, ICA without CFA removed, and uncorrected EEG). In a first instance, a Mann-Whitney U test was computed to compare the effect of each filtering method on the number of epochs exceeding amplitude threshold. This was followed by linear mixed model (LMM) analyses to determine which independent variables derived from the literature significantly affected the derived HEP values at the aforementioned EEG leads (figure 3). LMMs were performed as the dataset has a nested hierarchical structure with within-subject dependency. LMMs allow for explicit modelling of interacting terms while accounting for random effects including repeated measures, individual differences and unbalanced data. This approach aimed to preserve statistical power by incorporating these complexities into the analysis.

### 2.5. Artefact Correction methods

EEG is comprised of electrical potentials originating from various sources (Ungureanu et al., 2004). Artefacts from physiological sources (cardiac field artefact, pulse artefact, myogenic artefact) and non-physiological sources (electrode pops, movement) generate electrical activity that contaminate this EEG data. ICA and ASR are two distinct source-separation methods. These are effective techniques that are widely used to remove signal components related to non-brain activity and improve the signal quality of EEG (Plechawska-Wójcik et al., 2023). ICA and ASR were applied iteratively on the EEG data to subsequently compare their effect on artefact removal and HEP signal derivation.

#### 2.5.1. Independent component analysis

ICA seeks to find and disentangle statistically independent components that are linearly mixed across multiple sensors. Data is assumed to be non-gaussian. During signal processing, ICA coefficients are computed and are used to determine that spatial contribution on the signal component (Aljobouri, 2023; Carvalho, 2023). Components that appear artefactual are identified and removed, and the EEG signal is then reconstructed without the artefact (Ungureanu et al., 2004). ICA may require some initial manual intervention to categorise components as artefactual or EEG. In this review, ICA was applied using functions from the mne-iclabel package (version 0.6.0) for MNE (Li et al., 2022), and then artefactual components were identified using mne-icalabel. The ICA then fitted to the original EEG signal, at the pre-processing stage after picking clean sections of EEG from the original EEG data (figure 3) (Li et al., 2022).

#### 2.5.2. Artefact Subspace Reconstruction

ASR is a newer and more automated method which models and removes subspaces created by artefacts. The ASR process can be divided into three steps (Chang et al., 2019). First, reference data is extracted with ASR automatically detecting clean segments of raw EEG data based on the distribution of signal variance (Chang et al., 2019). The second step involves establishing thresholds for determining artefact components. Lastly, it eliminates artefact components and reconstructs cleaned data. One of the most important user-defined parameters in ASR is the cut-off parameter denoted as ‘***k’*** (Chang et al., 2019). This user defined parameter describes the rejection thresholds in units of standard deviation, i.e. ‘***k’*** is the parameter of the ASR algorithm regulating the threshold above which components would be identified as artifacts. Chang et al. (2019) demonstrated that the value of ‘k’ affects the performance of ASR for removing artefacts. The present study employed k values of 5, 10, and 20. In our analysis we applied ASR using the ASRpy package (Kothe and Jung 2016), an implementation of ASR in the python programming language.

## 3. Results

### 3.1. Literature scoping review – gaps and inconsistencies in HEP processing methods

Analysis of the HEP literature using the variables detailed previously, demonstrated a high level of heterogeneity in terms of the measures reported when deriving HEP in both EEG and MEG studies (table 4, table 5). These results highlighted the lack of consistency in reporting variables used for HEP derivation. The variables the least reported in published EEG studies were the epoch amplitude rejection threshold (80.2%), the number of samples (epochs) used for analysis (92%) and the minimum RR interval (97%) (table 4). Similar results were noted for MEG results where no studies reported on the epoch amplitude rejection value, and the number of samples used for analysis (table 5).

**Table 4.** The number of studies and the corresponding percentage where variables were unreported based on the 101 extracted EEG studies.

| Unreported variables | Number of studies | Percentage (%) |
| --- | --- | --- |
| Low-pass and/or high-pass filters | 25 | 25 |
| Epoch window | 5 | 5 |
| Epochs relative to the R wave or T wave | 3 | 3 |
| Epoch amplitude rejection threshold ($\mu\text{V}$) | 81 | 80 |
| HEP windows | 16 | 16 |
| Number of samples used for analysis | 93 | 92 |
| Baseline correction | 52 | 51 |
| Minimum RR interval | 98 | 97 |
| CFA correction | 38 | 38 |
| Electrodes used unreported | 28 | 28 |

**Table 5.** The number of studies and the corresponding percentage where variables were unreported based on the 10 extracted MEG studies.

| Unreported variables | Number of studies | Percentage (%) |
| --- | --- | --- |
| Low-pass and high-pass filters | 2 | 20 |
| Epoch window | 1 | 10 |
| Epochs relative to the R wave or T wave | 3 | 30 |
| Epoch amplitude rejection threshold (T) | 10 | 100 |
| HEP windows | 6 | 60 |
| Number of samples used for analysis | 10 | 100 |
| Baseline correction | 9 | 90 |
| Minimum RR interval | 9 | 90 |
| CFA correction | 1 | 10 |

### 3.2. Literature scoping review – heterogeneity in HEP parameter values

The literature demonstrated a significant level of heterogeneity in the values used for each parameter for HEP derivation (figure 4). This is further supported by the 108 unique combinations of preprocessing parameters (low-pass, high-pass, start and end time of epoch window, start and end time of HEP window, start and end time of baseline correction window, RR interval) extracted from the literature. Modal values may support the development of consensus values for HEP computation (table 6). Indeed, 26 studies (26%) out of 101 studies, used 30 Hz as the low-pass filter cut-off frequency (A) with a range from 15 Hz to 100 Hz. The two most common (modal values) high-pass cut-off frequencies (B) were 0.1 Hz and 0.5 Hz, used in 26 studies (26%). Fifty-two studies applied an epoch start time (C) at -200 ms and 49 studies (49%) utilised 600 ms as the epoch end time (D), in relation to the reference point utilised in the paper (R or T wave or unspecified). Two hundred milliseconds was identified as the modal value for the start of the HEP window (E) across 33 studies (33%). These studies showed an overall spread ranging from 0 ms to 550 ms, measured in relation to the R wave. Studies using the T wave as a reference point or those that were unspecified were excluded as these represented 4 of the total number of studies processed. Thirty-four studies (34%) employed 600 ms as the end of the HEP window (F) with values starting from 150 ms to 700 ms. Twenty-six studies (26%) used -200 ms for the start of the baseline correction window (G) with a range from -300 ms to 200 ms. For 44 studies (44%), 0 ms was the modal value for the end time of the baseline correction (H). Lastly, 5 studies (5%) used 50 µV as the epoch amplitude rejection threshold (I) with another 5 studies (5%) employing 100 µV as the threshold. The range of values spread from 30 µV to 300 µV.

**Figure 4.**
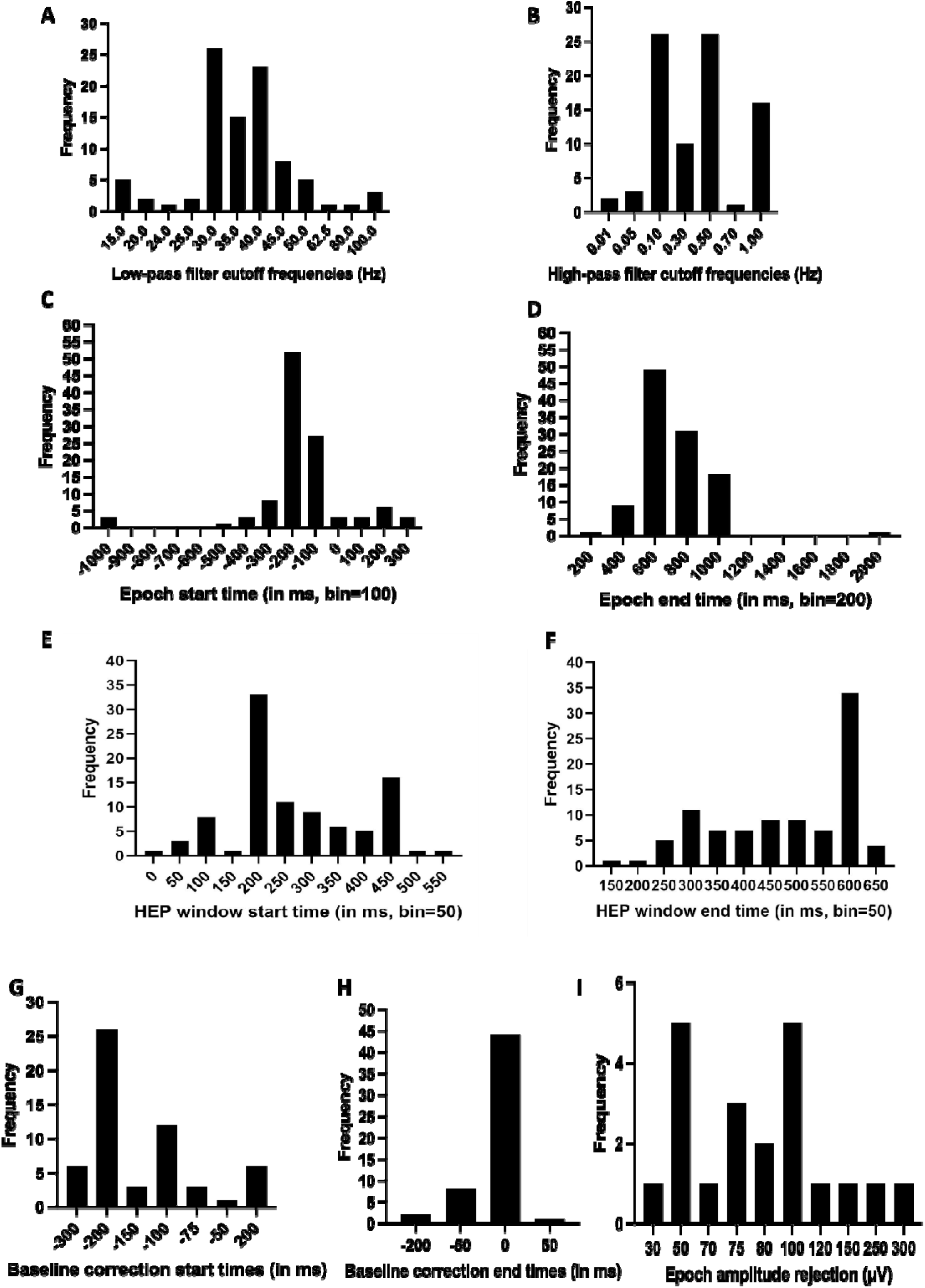
Values obtained from the literature used for each parameter (low pass (A) and high pass (B) filter cut-off frequencies, epoch start time (C) and end time (D), HEP window start time (E) and end time (F), baseline correction start time (G) and end time (H), epoch amplitude rejection (I) to derive HEP.

**Table 6:** Modal values identified in the literature and the number of studies employing these values. Where there were two modal values found for the same variable, the chosen modal value for that HEP variable has been highlighted in bold.

| HEP variable | Modal value | Number of studies | Percentage (%) |
| --- | --- | --- | --- |
| Low-pass filter | 30 Hz | 26 | 26 |
| High-pass filter | **0.1 Hz** | 26 | 26 |
| High-pass filter | 0.5 Hz | 26 | 26 |
| Epoch window start time | -200 ms | 52 | 51 |
| Epoch window end time | 600 ms | 49 | 49 |
| Start of the HEP window | 200 ms | 33 | 33 |
| End of the HEP window | 600 ms | 34 | 34 |
| Start of baseline correction window | -200 ms | 26 | 26 |
| End of baseline correction window | 0 ms | 44 | 44 |
| Epoch amplitude rejection threshold | 50 $\mu\text{V}$ | 5 | 5 |
| Epoch amplitude rejection threshold | **100 $\mu\text{V}$** | 5 | 5 |

### 3.3. Sensitivity analysis of HEP processing parameters

To assess the impact of each processing parameter (minimum RR, start and end times of the HEP window and start and end times of the baseline correction window) involved in deriving HEP, EEG data was epoched according to different parameter values extracted from the literature (figure 3). Figure 5 describes the relation between the number of epochs (in percentage) below the minimum RR threshold. As the RR interval increased, more epochs were dropped for further analysis as they did not meet the RR threshold, i.e. those epochs were too short. Epochs where the time until the next R wave was less than the minimum RR threshold may increase the risk of CFA affecting the HEP. Figure 6 describes the relation between the percentage of epochs exceeding the amplitude threshold and the epoch amplitude rejection threshold. The percentage of epochs excluded decreased rapidly when the maximum epoch amplitude was below 100 µV, but above this value remained relatively constant at 40 epochs (5%). When the amplitude threshold is varied, there is a clear threshold effect; amplitudes below 100 µV lead to an exponential increase in the number of epochs rejected. Conversely, above 100 µV, there is little change in the number of epochs rejected. In wakefullness, EEG is mostly under 100 µV. Therefore, there is not a significant change in the number of epochs exceeding the amplitude rejection threshold passed 100 µV. Figure 7 represents the number of epochs exceeding amplitude threshold following the application of the different filtering techniques identified in the literature (ASR 5, ASR 10, ASR 20, ICA without CFA removed and uncorrected EEG) using the modal values defined for high pass filters (0.1 Hz), low pass filters (30 Hz), maximum epoch amplitude rejection (100 µV) and minimum RR interval (0.88 s). Overall, ASR-5, ASR-10, ASR-20 and ICA without CFA removed seem to yield similar results in terms of the reduction in epochs exceeding the amplitude threshold.

**Figure 5.**
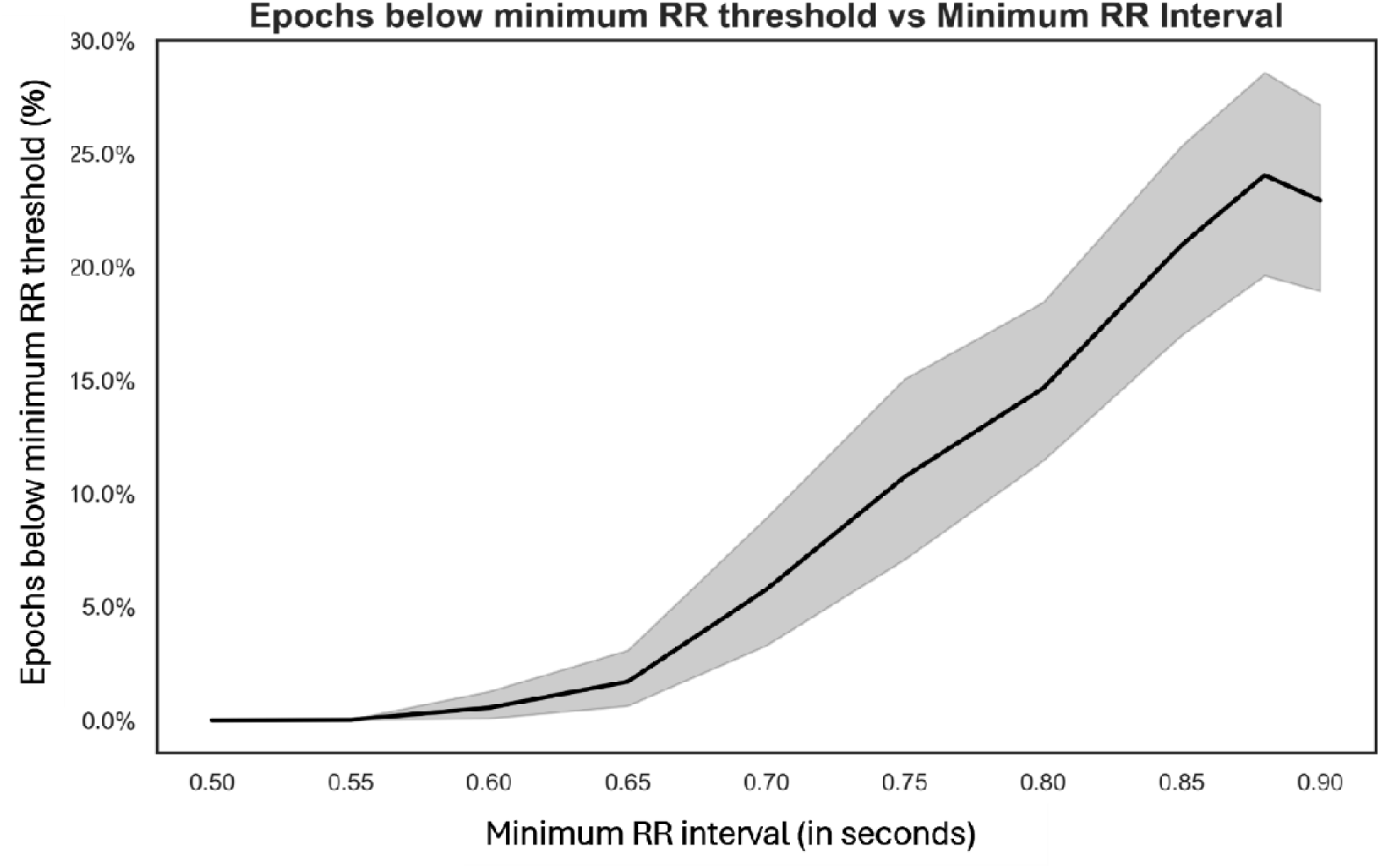
The number of epochs (represented in %) below minimum RR threshold according to the minimum RR interval (in milliseconds) across all filtering methods (ICA, ASR, and uncorrected EEG).

**Figure 6.**
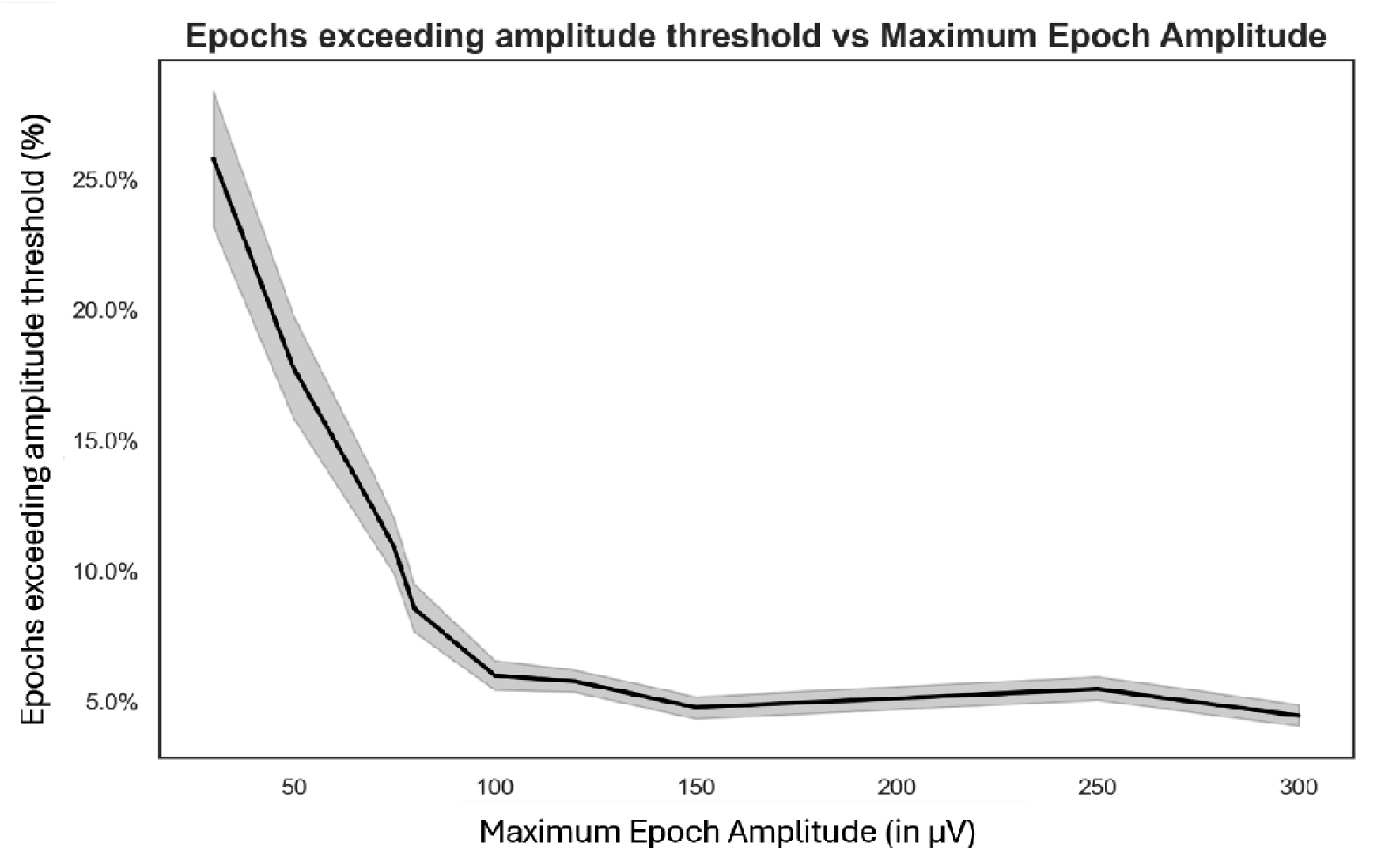
The number of epochs (represented in %) with too much artefact according to the maximum epoch amplitude (in µV) across all filtering methods (ICA, ASR, and uncorrected EEG).

**Figure 7.**
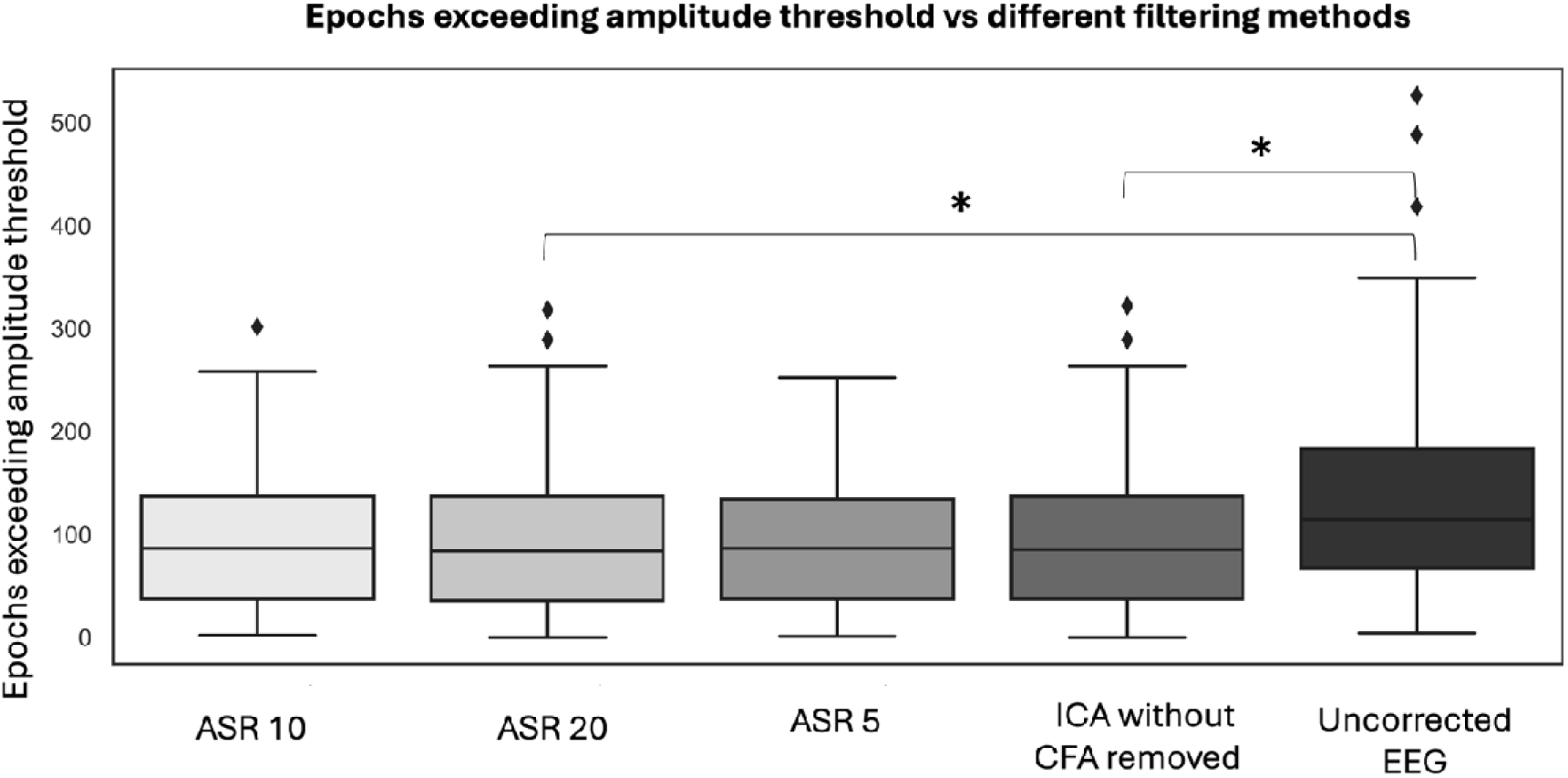
Box plot representing the number of epochs exceeding the amplitude threshold in relation to the different filtering parameters using the modal values defined for high pass filters (0.1 Hz), low pass filters (30 Hz), epoch amplitude rejection (100 µV) and minimum RR interval (0.88 seconds).

Figure 8 depicts the grand averages of EEG evoked potentials as voltage maps. This analysis was conducted to increase signal-to-noise ratio and reveal evoked potential morphology across different filtering methods. Each point on the voltage map represents the instantaneous voltage at that lead. Panel A represents grand averages of uncorrected EEG data which is noisy as expected. The concentration of signals in the time series of panel B (ICA without CFA removed) suggests that the ICA without CFA removal method could have effectively removed artefacts and noise. Panel E with a less aggressive ASR threshold of 20, seems to exhibit a similar signal to ICA without CFA removed. Grand averages in panel D with a more constraining ASR threshold (ASR-10) seem to contain more oscillations than other panels (i.e. ASR-5). Upon investigation, oscillations in ASR-10 appear to be resulting from specific individual event-related potential (ERPs) from the 97 EEG recordings included in this review. These ERPs seem to contain oscillations that skew the averaged EEG evoked potential. EEGs including these abnormal oscillations were not excluded from further statistical analysis. Observations are further supported with statistical analysis in 3.6.1.

**Figure 8.**
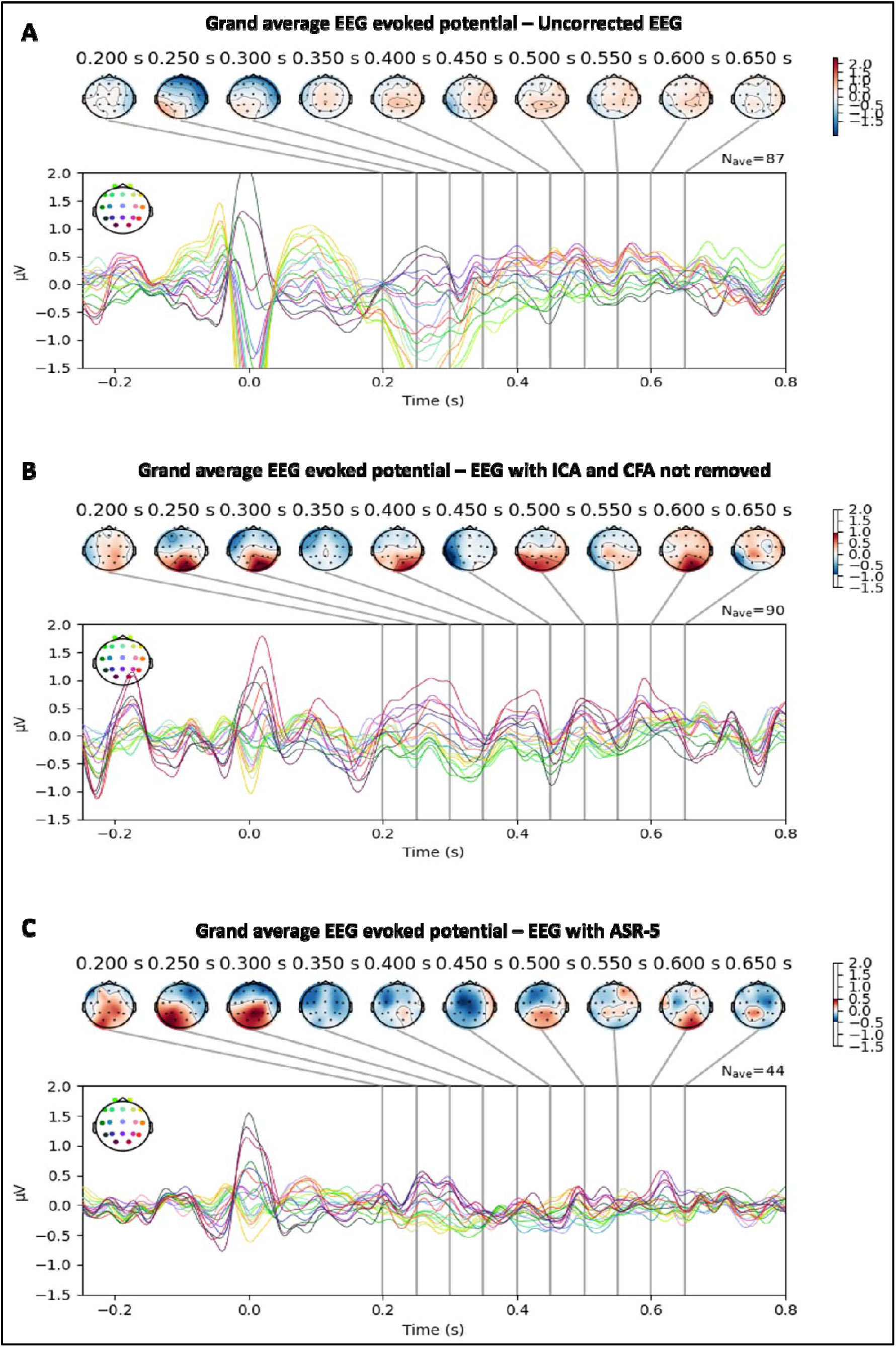
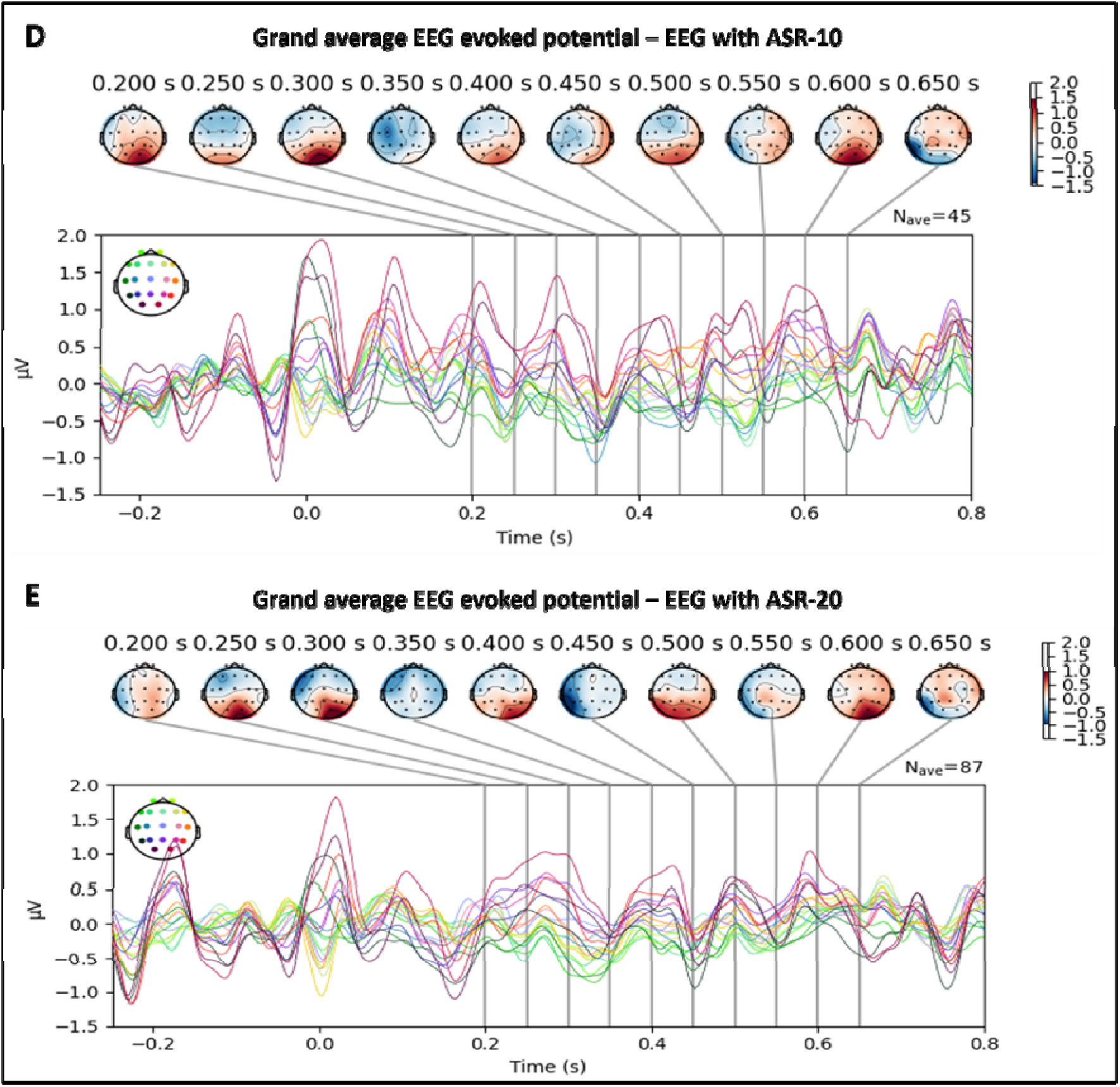
Grand average topographical voltage maps across filtering methods.

### 3.4. Sensitivity analysis: Mann-Whitney U test - effect of artefact correction methods on the number of epochs exceeding maximum epoch amplitude threshold

A Mann-Whitney U test (independent samples T-test) was conducted to assess the effect of different EEG filtering methods on the number of epochs exceeding the maximum epoch amplitude threshold (figure 7, figure 9). This non-parametric statistical test was chosen as the data was non-normal. After Bonferroni correction, α was set at 0.01. While there was no significant difference between the different artefact correction methods, there was a significant difference between the artefact correction methods and uncorrected EEG, particularly between ASR-20 and uncorrected EEG (p<0.001) and ICA without CFA removed and uncorrected EEG (p=0.002) after Bonferroni correction.

**Figure 9:**
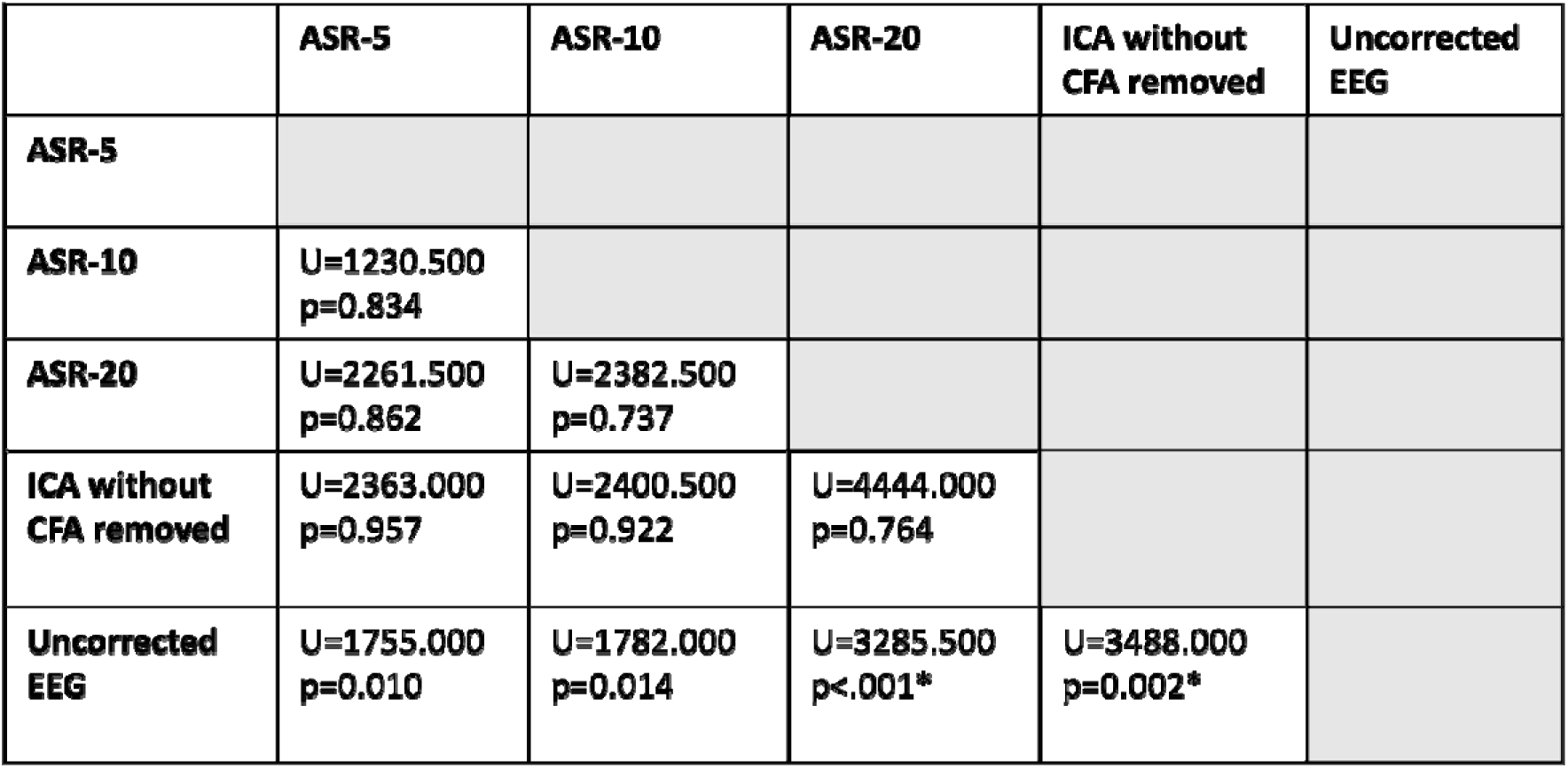
Matrix representation of Mann-Whitney U test. Results illustrate the effect of the filtering methods explored on the number of epochs exceeding the maximum epoch amplitude threshold. Bonferroni correction was set at α = 0.01. All other variables were kept at their modal values (high pass filter – 0.1 Hz, low pass filter – 30 Hz, epoch amplitude rejection – 100 µV, minimum RR interval – 0.88 seconds).

### 3.5. Influence of artefact correction methods and preprocessing parameters on HEP variability: heatmap analysis

Heatmaps, in figure 10 and 11, represent averaged HEP values and standard deviations in HEP, in each artefact correction method when only one processing parameter is being varied while others are kept at their modal value.

**Figure 10:**
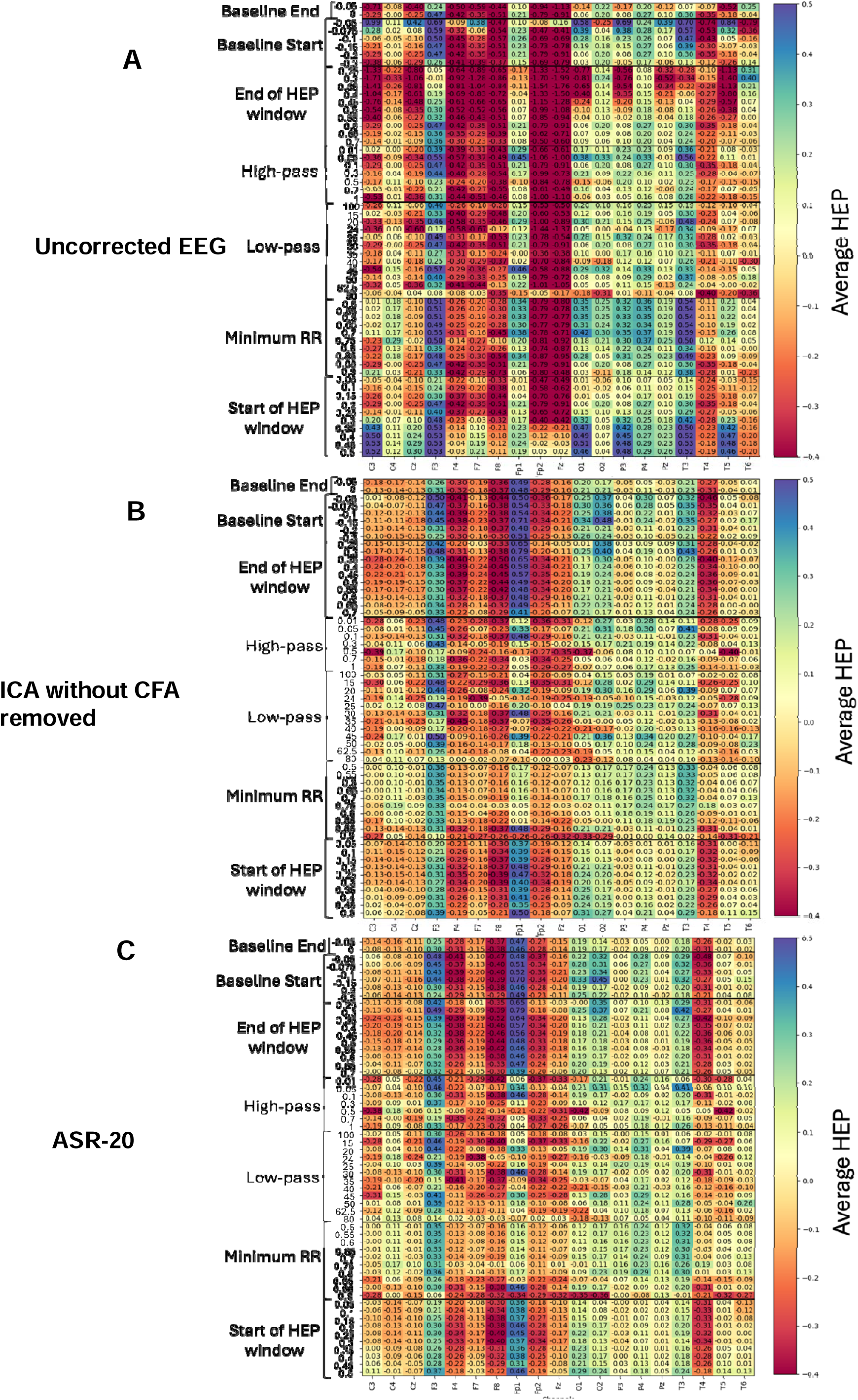
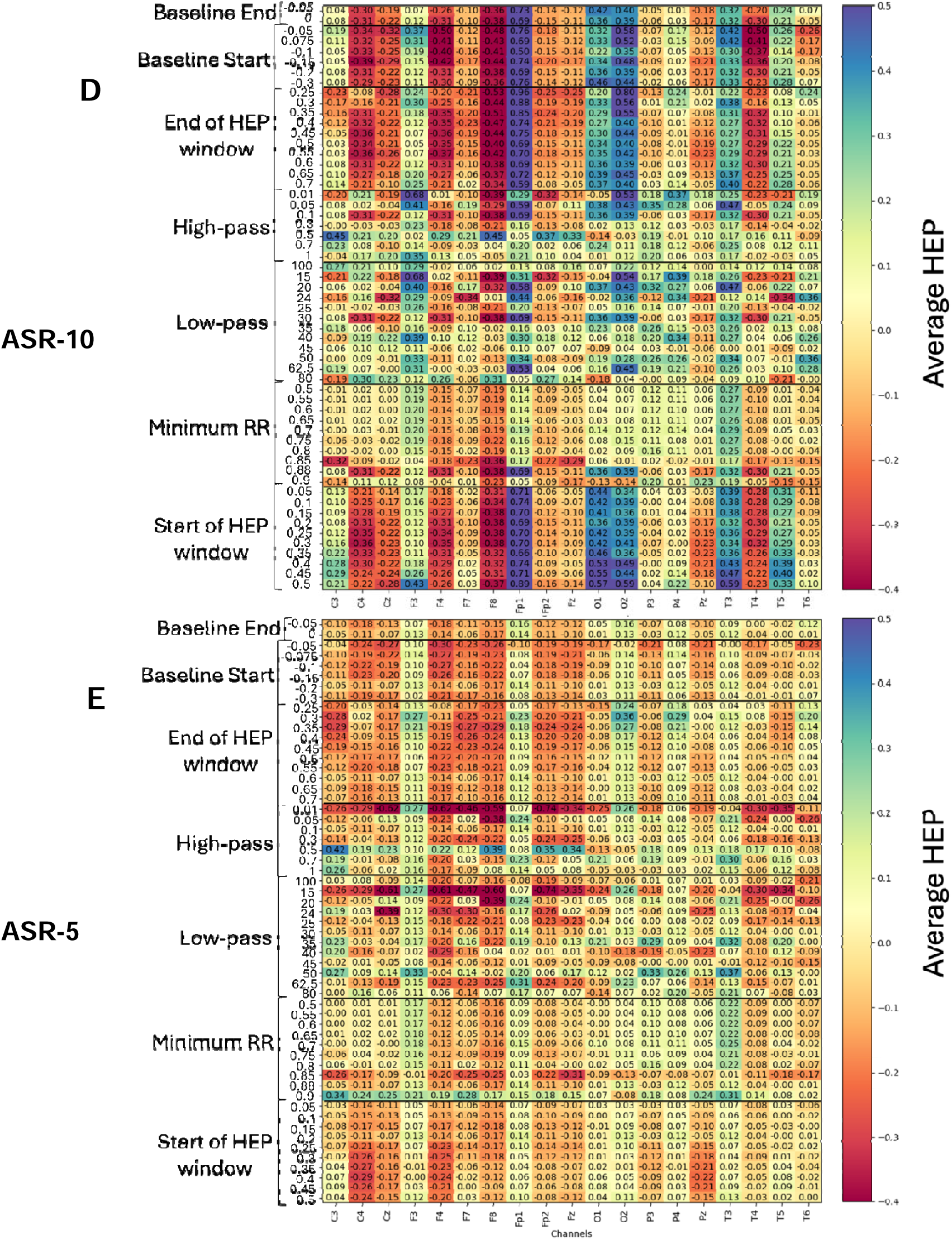
Heatmaps representing average HEP values for different artefact correction methods and processing settings. Average HEP values are shown for varying processing parameters (rows) across EEG channels (columns). Artefact correction methods include Independent Component Analysis (ICA), Artefact Subspace Reconstruction (ASR) with thresholds of 5, 10, and 20, and uncorrected raw EEG data. Each processing parameter (baseline start/end, HEP window start/end, high-pass/low-pass filters, and minimum R-R interval) is varied independently while others are held at their modal values. Colour intensity represents the magnitude of average HEP values.

In figure 10, across all values for start and end of baseline correction, there is high variability in HEP values, indicating high sensitivity of HEP to baseline correction windows. Across ICA without CFA removed, ASR-10, and ASR-20, HEP is consistently more positive in channel Fp1 compared to other conditions. However, in the uncorrected raw EEG, HEP appears visually more negative in certain channels (F4, F8, Fp2, Fz). This same pattern is found in the start and end of the HEP window, suggesting sensitivity of HEP features to the selected time window. The effect of minimum RR on HEP appears consistent across all artefact correction methods. In general, the patterns observed in ASR-20 and ICA without CFA appear to be similar, indicating they may be influenced in equivalent ways by processing parameters. ASR-5 appears to be visually very different to uncorrected EEG. Overall, HEP changes considerably and can become more negative or positive when going from one value to another within each processing parameter.

In figure 11, standard deviations appear to be higher in uncorrected EEG compared to all other artefact correction methods. Standard deviations across all channels and processing settings in ASR-5 are noticeably lower compared to ASR-10, ASR-20 and ICA without CFA removed. Channel Fp1 demonstrates high standard deviations for start and end of the HEP window and start of the baseline correction window in ICA without CFA removed, ASR-5, ASR-10 and ASR-20. Artefact seems to be efficiently removed by low-pass and high-pass filters and by changing the minimum RR, as demonstrated by the low standard deviations in those processing parameters. More specifically, minimum RR seems to have a noticeable effect on reducing HEP variability. However, it appears that minimum RR exhibits a threshold effect, where the specific value of the minimum RR interval becomes irrelevant if it exceeds a certain threshold.

**Figure 11:**
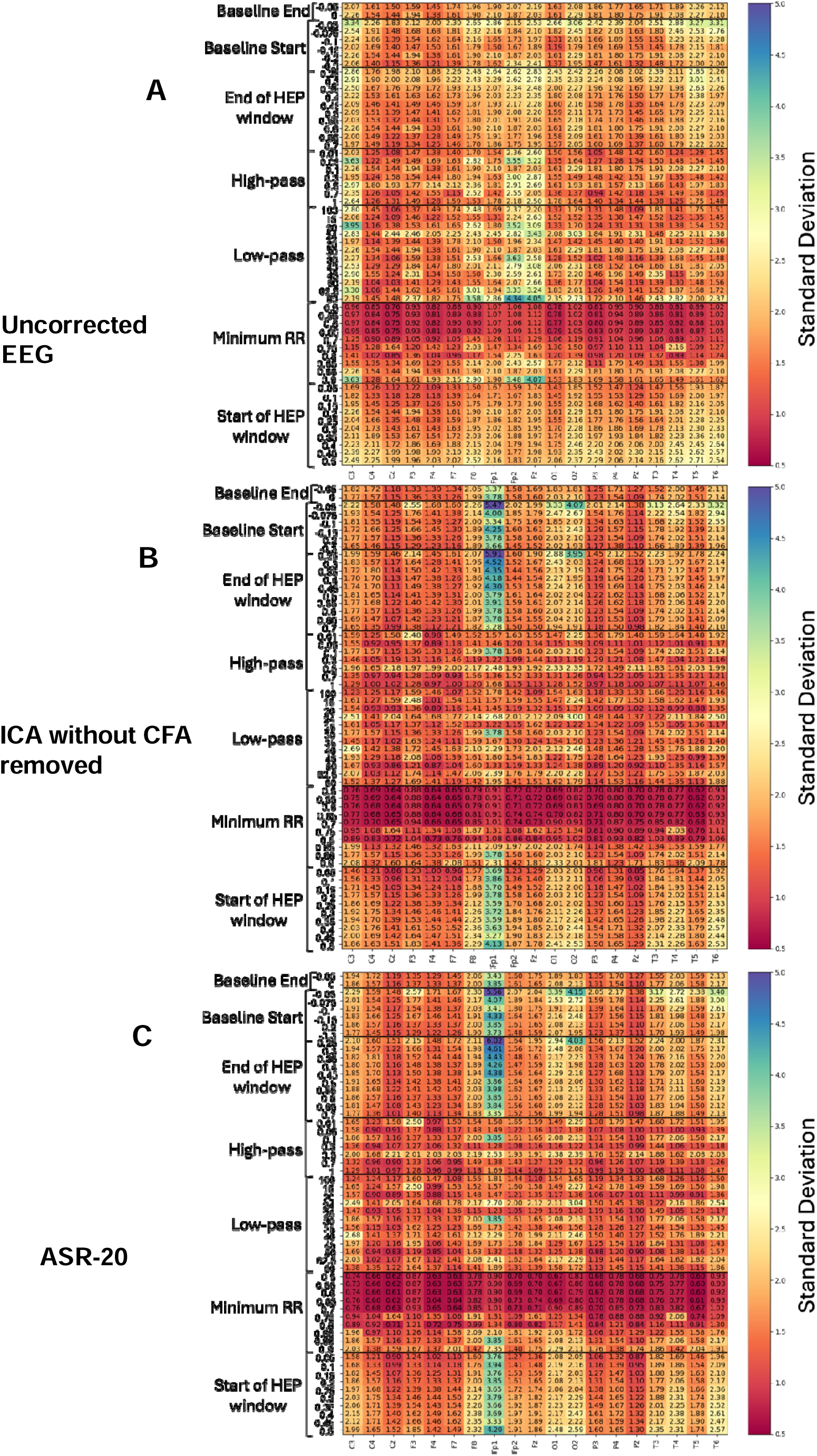
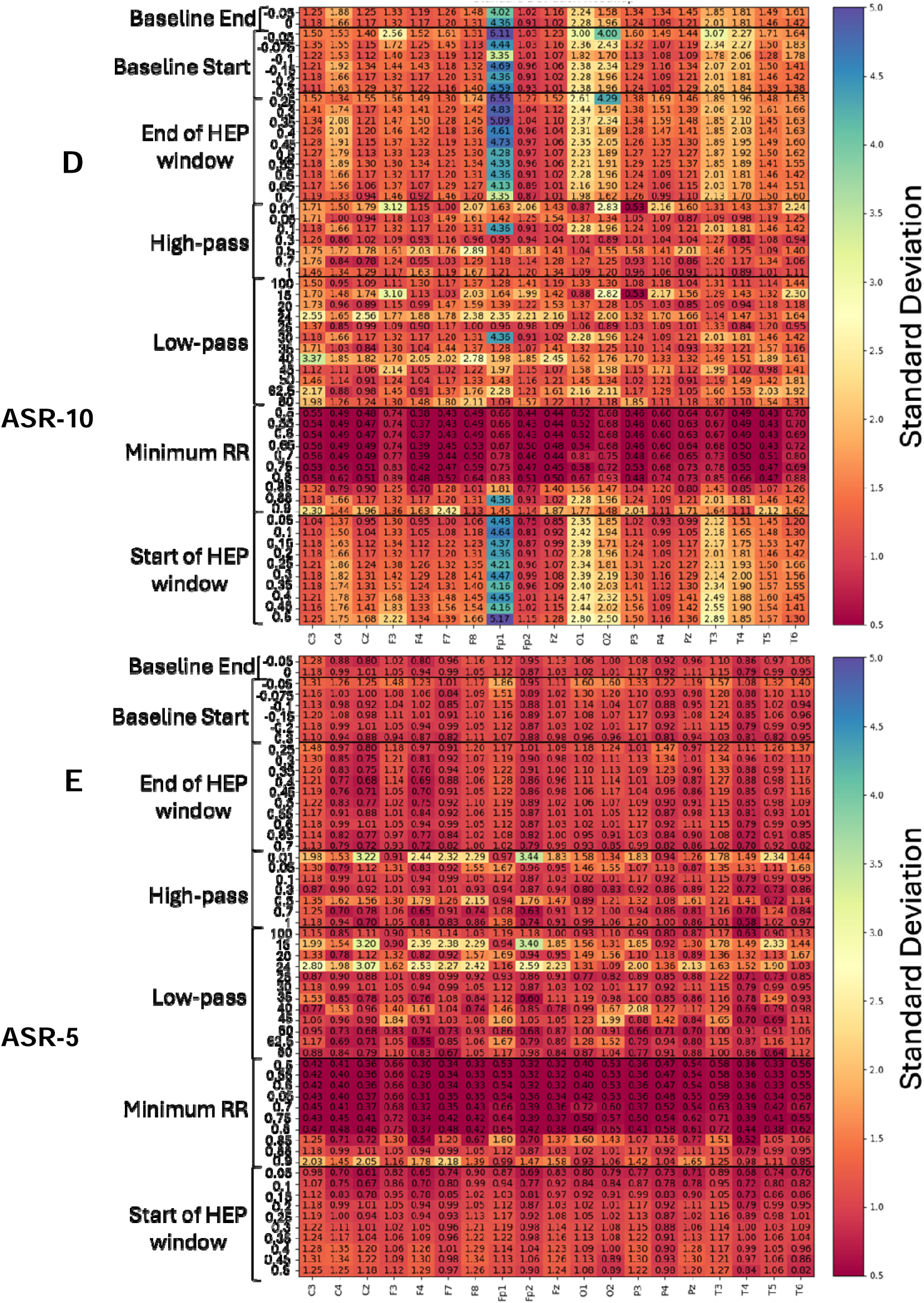
Heatmaps representing standard deviation in HEP values for different artefact correction methods and processing settings. Heatmaps show the standard deviation of HEP values for different artefact correction methods (ICA without CFA removed, ASR-5, ASR-10, ASR-20, and uncorrected EEG) and processing parameters (rows). Each processing parameter is varied independently, while others remain constant. Higher standard deviation (yellow to blue colours) indicates greater variability across conditions.

### 3.6. Linear mixed model (LMM) analysis on artefact correction methods (ICA, ASR-5, ASR-10, ASR-20, uncorrected EEG) and processing settings

To further explore differences in HEP related to parameter choices, LMMs were conducted by considering artefact removal methods, processing settings and channels. Statistical analysis was performed in Python (version 3.11) using the statsmodels.formula.api package (version 0.14.4) and pingouin package (version 0.5.5) for calculating estimated marginal means (EMMs).

#### 3.6.1. LMM analysis on pairwise comparisons between artefact correction methods and channels

To assess the impact and interaction of artefact correction methods and channels on HEP values, an LMM was conducted with artefact removal methods and channels as the fixed effects and ICA without CFA removed as the reference level (gold standard used in the literature). Random effects in the model included: subject, channels, low-pass filter, high-pass filter, minimum RR threshold, maximum epoch amplitude, start and end of HEP window and start and end of baseline correction window. The model included subject as a primary grouping factor for random effects. Results in table 7 demonstrate that ASR-5, ASR-10 and uncorrected EEG relative to ICA without CFA removed have a significant effect on HEP values. Indeed, HEP values when using ASR-5 (β=0.082, p=0.010) and ASR-10 (β=0.119, p<0.001) are significantly higher than when using ICA without CFA removed. HEP values derived from uncorrected EEG are significantly lower than those derived from ICA corrected EEG (β=-0.118, p<0.001). HEP values were similar for ASR-20 corrected EEG compared to ICA corrected (β=0.019, p=0.448).

**Table 7:** Linear Mixed Model Results - pairwise comparisons of artefact correction methods on HEP Values. S.E.: standard error; Z val.: Z value; P-unc: P uncorrected.

|  | **Estimate** | **S.E.** | **Z val.** | **P-unc** |
| --- | --- | --- | --- | --- |
| **INTERCEPT** | 0.006 | 0.157 | 0.037 | 0.971 |
| **ASR-5 - ICA** | 0.082 | 0.032 | 2.561 | 0.010 |
| **ASR-10 - ICA** | 0.119 | 0.032 | 3.747 | <0.001 |
| **ASR-20 - ICA** | 0.019 | 0.025 | 0.758 | 0.448 |
| **Uncorrected - ICA** | -0.118 | 0.025 | -4.742 | <0.001 |

LMMs were followed by post hoc analysis using estimated marginal means (EMMs) (table 8). ASR-5 and ASR-10 appear to differ significantly (p<0.001***\****), with ASR-10 showing slightly higher effects. Significant differences are found between ASR-10 and ICA (p<0.001***\****), but with a very small effect size (Hedges’ g = 0.027). There are strong differences between ASR-10 and uncorrected data (p<0.001***\****). There are no significant differences between ASR-20 and ASR-5 (p = 0.053, Hedges’ g =0.013).

**Table 8:** Estimated marginal means post hoc analysis. Stars indicate significance after EMM. P-unc: p uncorrected, p-corr: p corrected.

|  | P-unc | P-corr | BF10 | Hedges' g |
| --- | --- | --- | --- | --- |
| **ASR-5 - ICA** | 0.010 | 0.0044* | 2.907 | -0.017 |
| **ASR-10 - ICA** | <0.001 | <0.001* | 3396.985 | 0.027 |
| **ASR-20 - ICA** | 0.448 | 1 | 0.006 | -0.003 |
| **Uncorrected - ICA** | <0.001 | <0.001* | $5.42 \times 10^{30}$ | 0.054 |
| **ASR-20 - ASR-10** | <0.001 | <0.001* | $6.47 \times 10^{04}$ | 0.030 |
| **ASR-5 - ASR-10** | <0.001 | <0.001* | $3.15 \times 10^{12}$ | 0.051 |
| **Uncorrected – ASR-10** | <0.001 | <0.001* | $2.33 \times 10^{50}$ | 0.079 |
| **ASR-5 – ASR-20** | 0.005 | 0.053 | 0.299 | 0.013 |
| **Uncorrected – ASR-20** | <0.001 | <0.001* | $1.06 \times 10^{27}$ | 0.052 |
| **ASR-5 – uncorrected** | <0.001 | <0.001* | $8.98 \times 10^{16}$ | 0.043 |

Pairwise comparisons investigating interaction effects between artefact settings and channels are included in the supplementary material (see supplementary material).

#### 3.6.2. LMM analysis on processing settings

To investigate the impact and interaction of processing parameters and channels on HEP values, an LMM was conducted with all processing parameters and channels as the fixed effects. Random effects in the model included: patients, channels, and artefact correction methods. The model included patients as a primary grouping factor for random effects.

Results in table 9 demonstrate the minimum RR (β<0.001, p<0.001), maximum epoch amplitude (β<0.001, p<0.001), start and end of baseline correction (β<0.001, p<0.001) and start and end of HEP window (β<0.001, p<0.001) have a significant effect on HEP. HEP values do not appear to be impacted by low-pass (β<0.001, p=0.136) or high-pass filters (β<0.001, p=0.875) although heatmaps in figure 10 appear to indicate that HEP values change considerably with different values for low-pass and high-pass filters.

**Table 9:** Linear Mixed Model Results – effect of processing settings on HEP values. S.E. – standard error; Z val. – Z value.

|  | **Estimate** | **S.E.** | **Z val.** | **p** |
| --- | --- | --- | --- | --- |
| **Intercept** | 0.141 | 0.067 | -2.117 | 0.034 |
| **Low-pass** | <0.001 | 0.000 | 1.490 | 0.136 |
| **High-pass** | <0.001 | 0.000 | -0.157 | 0.875 |
| **Minimum RR** | <0.001 | 0.000 | -10.127 | <0.001 |
| **Maximum epoch amplitude** | <0.001 | 0.000 | 3.606 | <0.001 |
| **Baseline start** | <0.001 | 0.000 | 3.711 | <0.001 |
| **Baseline end** | 0.001 | 0.000 | 3.506 | <0.001 |
| **Start of HEP window** | <0.001 | 0.000 | 10.584 | <0.001 |
| **End of HEP window** | <0.001 | 0.000 | 15.878 | <0.001 |

LMMs were followed by post hoc analysis using estimated marginal means (EMMs) to investigate the relation between the values of each processing parameter and HEP. These results are included in the supplementary material (see Supplementary material).

## 4. Discussion

Results from the literature scoping review demonstrated a significant level of heterogeneity in the values used for HEP derivation and lack of consistency in terms of variables reported in studies. This demonstrates the fundamental need for a processing pipeline to derive HEP to ensure replicability and consensus in this growing field.

As expected, increasing the allowed RR interval increases the number of epochs that meet the criteria (figure 5). Epochs shorter than the minimum RR threshold could cause the infiltration of cardiac field artefacts which would impact HEP computation and may lead to inaccurate results. Epochs containing activity with amplitudes exceeding the amplitude threshold value are thought likely to do so due to artefact, rather than EEG activity, and higher amplitudes are more likely to skew averaged evoked responses (figure 6). Investigating the impact of artefact correction methods on the number of epochs exceeding amplitude threshold, demonstrated that all artefact correction methods remove epochs with large amplitude artefacts from EEG data (figure 7). P-values were not significant after Bonferroni correction for ASR-5 and ASR-10 versus uncorrected EEG. However, ASR-20 and ICA without CFA removed showed significant results for removing artefact from EEG in comparison to uncorrected EEG (figure 9).

Topographies of the evoked EEG potentials (figure 8) and LMM analyses (tables 7 and 8) demonstrated that artefact correction methods significantly affect HEP. As expected, ICA and ASR seem to clearly affect HEP versus uncorrected EEG (table 7), supporting the use of artefact rejection in deriving HEP. The choice of artefact correction method significantly influences HEP values, as evidenced by statistical significance in the LMM analysis. However, although differences are statistically significant, the effect size are mostly small to negligible, suggesting that while artefact correction impacts HEP values, the magnitude of the impact is limited. This could be due to a limitation in the statistical method used which assumes a linear relationship between predictors and response variables. Based on the prior analyses, it seems these signal processing methods (ASR-20 and ICA) reduce the number of rejected epochs based on exceeding an amplitude threshold (figure 9, table 7). This may suggest that ASR using a ‘***k’*** value threshold of 20 is as performant as ICA without CFA removed (table 8). Indeed, Plechawska-Wójcik et al. (2023) suggested the optimal ASR parameter to be between 20 and 30. However, a delicate balance must be struck to remove artefacts efficiently without removing or aberrantly altering the EEG signal. Overall, this is an exciting finding as ICA has been predominantly used in the literature until now. Given that ASR can be utilised concurrently with EEG acquisition, its potential application in future wearable technologies might be possible. ICA and ASR-20 could then be the optimal artefact correction methods to use in HEP analysis.

LMM analysis (table 9) demonstrated that the RR interval, the maximum epoch amplitude, the start and end of the HEP window and the start of the baseline correction window significantly affect HEP as described previously. From the heatmap (figure 11), minimum RR appears to have a threshold effect on reducing HEP variability. The specific value of the RR interval becomes irrelevant as long as it exceeds a certain threshold. The RR interval represents the interval in which HEP is found and comprises of one heartbeat. The minimum RR is therefore the smallest interval in which a single HEP amplitude can be found. If this interval is too short, there is a greater risk that the HEP value derived would be contaminated by the CFA, which can be an order of magnitude greater than the HEP signal. Given this significant effect, it is concerning that 97% of EEG studies do not report either whether a minimum RR interval has been used and, if so, at what value. The RR interval should be carefully considered during EEG analysis. HEP derivation in participants with tachycardia is at risk of finding an inflated HEP value. The start and end of the HEP window are critical for deriving HEP and minimising contamination by the CFA. As demonstrated by the scoping review, about 16% of studies do not report on this value, despite how critical it is to HEP analysis. Similarly, approximately 52% of studies do not explicitly state the baseline correction window, which this review has demonstrated to affect the HEP. As investigated, CFA correction is crucial for a reliable HEP value. The literature revealed that ≈38% of studies do not report whether CFA correction was performed or not. These results highlight the lack of consistency within the field. Hence, particular care should be given to these variables when acquiring and deriving HEP to ensure consistency and reproducibility of analysis. Parameters used for HEP extraction should be reported in publications, emphasising the need for standardised methods to enhance study comparison and reproducibility and establish a gold standard in the field.

## 5. Conclusion

The present study suggests that there is vast heterogeneity and variability in the methods used to derive HEP. There is a clear inconsistency in the use of parameters to derive HEP as well as reporting these in the literature. A significant number of parameter choices seem to affect the HEP value. Consequently, the level of reproducibility and reported results appears to be variable. Artefact correction methods appear to remove artefact in general and epoch rejection seems to be equivalent between ICA and ASR-20. This could indicate that ICA and ASR-20 are the optimal methods, or at least consistent methods, to use when deriving HEP. Certain processing parameters do influence HEP significantly. These include the RR interval, the maximum epoch amplitude, the start and end of the HEP window and the start of the baseline correction window significantly affect HEP and should be carefully considered during HEP derivation. However, it is hard to understand the effect these parameters have on HEP in terms of direction and magnitude. Values for HEP extraction should be reported in publications, to enhance study comparison and reproducibility and eventually establish a gold standard in the field. Pipelines used to derive HEP should be made available on repositories such as GitHub. While there is no “ground truth” for HEP, this review has aimed to investigate how HEP extraction parameters influence and affect the resulting HEP. Ultimately, to develop robust methods for HEP derivation, it is essential to establish a “ground truth” for HEP with a clear consensus on CFA removal. Such standardisation would provide a reliable foundation for advancing research and enhancing the precision and accuracy of HEP. This review serves as a basis for further work in this matter.

## Data and Code Availability

The data underlying this article are available in The Temple University Hospital EEG data (TUH EEG) corpus, at https://doi.org/10.34944/dspace/5136. The code used for analysis is publicly available on GitHub at https://github.com/RaniaImanV/HEP-PARAMs.git.

## Author contributions/acknowledgments

All authors were involved in the conceptualisation, methodology, review and editing of the drafts and supervision (M.Y., S.N.G., D.C.). R.I.V. and R.K. were involved in the investigation, software development (primarily R.K.), and formal analysis. R.I.V. created the visualisation and wrote the original draft.

## Ethics statement

this study did not involve testing on humans or animals.

## References

1. Aljobouri, H. K. (2023). Independent component analysis with functional neuroscience data analysis. Journal of Biomedical Physics & Engineering, 13(2), 169–180. 10.31661/jbpe.v0i0.2111-1436

2. Arnau, S., Sharifian, F., Wascher, E., & Larra, M. F. (2023). Removing the cardiac field artifact from the EEG using neural network regression. Psychophysiology, 60(10). 10.1111/psyp.14323

3. Babo-Rebelo, M., Richter, C. G., & Tallon-Baudry, C. (2016a). Neural responses to heartbeats in the default network encode the self in spontaneous thoughts. Journal of Neuroscience, 36(29), 7829–7840. 10.1523/jneurosci.0262-16.2016

4. Baranauskas, M., Grabauskaitė, A., & Griškova-Bulanova, I. (2017). Brain responses and self-reported indices of interoception: Heartbeat evoked potentials are inversely associated with worrying about body sensations. Physiology & Behavior, 180, 1–7. 10.1016/j.physbeh.2017.07.032

5. Canales-Johnson, A., Silva, C., Huepe, D., Rivera-Rei, A., Noreika, V., Garcia, M. D., Silva, W., Ciraolo, C., Vaucheret, E., Sedeno, L., Couto, B., Kargieman, L., Baglivo, F., Sigman, M., Chennu, S., Ibanez, A., Rodriguez, E., & Bekinschtein, T. A. (2015). Auditory feedback differentially modulates behavioral and neural markers of objective and subjective performance when tapping to your heartbeat. Cerebral Cortex, 25(12), 4490–4503. 10.1093/cercor/bhv076

6. Chang, C.-Y., Hsu, S.-H., Pion-Tonachini, L., & Jung, T.-P. (2019). Evaluation of artifact subspace reconstruction for automatic artifact components removal in multi-channel EEG recordings. IEEE Transactions on Biomedical Engineering, 1–1. 10.1109/tbme.2019.2930186

7. Coll, M.-P., Penton, T. and Hobson, H. (2017). Important methodological issues regarding the use of transcranial magnetic stimulation to investigate interoceptive processing: a Comment on Pollatos et al. (2016). Philosophical Transactions of the Royal Society B: Biological Sciences, 372(1721), p.20160506. 10.1098/rstb.2016.0506

8. Coll, M.-P., Hobson, H., Bird, G., & Murphy, J. (2021). Systematic review and meta-analysis of the relationship between the heartbeat-evoked potential and interoception. Neuroscience & Biobehavioral Reviews, 122, 190–200. 10.1016/j.neubiorev.2020.12.012

9. Desmedt, O., Olivier Luminet Walentynowicz, M. and Corneille, O. (2023). The new measures of interoceptive accuracy: A systematic review and assessment. Neuroscience & Biobehavioral Reviews, 153, pp.105388–105388. 10.1016/j.neubiorev.2023.105388

10. Elkommos, S., Martín-López, D., Koreki, A., Jolliffe, C., Mula, M., Critchley, H. D., Edwards, M. J., Garfinkel, S. N., Richardson, M., & Yogarajah, M. (2022). Attenuated heart-brain integration predicts functional non-epileptic seizures. Journal of Neurology, Neurosurgery, and Psychiatry, 93(12), e3.7–e3.7. 10.1136/jnnp-2022-bnpa.15

11. Elkommos, S., Martín-López, D., Koreki, A., Jolliffe, C., Kandasamy, R., Mula, M., Critchley, H. D., Edwards, M., Garfinkel, S. N., Richardson, M., & Yogarajah, M. (2023). Changes in the heartbeat-evoked potential are associated with functional seizures. Journal of Neurology, Neurosurgery, and Psychiatry, 94(9), 769–775. 10.1136/jnnp-2022-330167

12. Flasbeck, V., Popkirov, S., Ebert, A., & Brüne, M. (2020). Altered interoception in patients with borderline personality disorder: A study using heartbeat-evoked potentials. Borderline Personality Disorder and Emotion Dysregulation, 7(1). 10.1186/s40479-020-00139-1

13. Garfinkel, S. N., & Critchley, H. D. (2016). Threat and the body: How the heart supports fear processing. Trends in Cognitive Sciences, 20(1), 34–46. 10.1016/j.tics.2015.10.005

14. Gray, M. A., Taggart, P., Sutton, P. M., Groves, D., Holdright, D. R., Bradbury, D., Brull, D., & Critchley, H. D. (2007). A cortical potential reflecting cardiac function. Proceedings of the National Academy of Sciences, 104(16), 6818–6823. 10.1073/pnas.0609509104

15. Hodossy, L., Ainley, V. and Tsakiris, M. (2021). How do we relate to our heart? Neurobehavioral differences across three types of engagement with cardiac interoception. Biological Psychology, 165, p.108198. 10.1016/j.biopsycho.2021.108198

16. Jones, G. E., Leonberger, T. F., Rouse, C. H., Caldwell, J. A., & Jones, K. R. (1986). Preliminary data exploring the presence of an evoked potential associated with cardiac visceral activity. Psychophysiology, 23, 445.

17. Kamp, S.-M., Schulz, A., Forester, G., & Domes, G. (2021). Older adults show a higher heartbeat-evoked potential than young adults and a negative association with everyday metacognition. Brain Research, 147238. 10.1016/j.brainres.2020.147238

18. Kern, M., Ad Aertsen, Schulze-Bonhage, A., & Ball, T. (2013). Heart cycle-related effects on event-related potentials, spectral power changes, and connectivity patterns in the human ECoG. NeuroImage, 81, 178–190. 10.1016/j.neuroimage.2013.05.042

19. Khalsa, S. S., Rudrauf, D., Feinstein, J. S., & Tranel, D. (2009). The pathways of interoceptive awareness. Nature Neuroscience, 12(12), 1494–1496. 10.1038/nn.2411

20. Khalsa, S. S., Adolphs, R., Cameron, O. G., Critchley, H. D., Davenport, P. W., Feinstein, J. S., Feusner, J. D., Garfinkel, S. N., Lane, R. D., Mehling, W. E., Meuret, A. E., Nemeroff, C. B., Oppenheimer, S., Petzschner, F. H., Pollatos, O., Rhudy, J. L., Schramm, L. P., Simmons, W. K., Stein, M. B., & Stephan, K. E. (2018). Interoception and mental health: A roadmap. Biological Psychiatry: Cognitive Neuroscience and Neuroimaging, 3(6), 501–513. 10.1016/j.bpsc.2017.12.004

21. Kim, K. J., Ramiro Diaz, J., Iddings, J. A., & Filosa, J. A. (2016). Vasculo-neuronal coupling: Retrograde vascular communication to brain neurons. Journal of Neuroscience, 36(47), 12624–12639. 10.1523/jneurosci.1300-16.2016

22. Kothe, C. A. E., & Jung, T. P. (2016). U.S. Patent Application No. 14/895,440. https://patents.google.com/patent/US20160113587A1/en

23. Kumral, D., Al, E., Cesnaite, E., Kornej, J., Sander, C., Hensch, T., Zeynalova, S., Tautenhahn, S., Hagendorf, A., Laufs, U., Wachter, R., Nikulin, V., & Villringer, A. (2022). Attenuation of the heartbeat-evoked potential in patients with atrial fibrillation. JACC: Clinical Electrophysiology, 8(10), 1219–1230. 10.1016/j.jacep.2022.06.019

24. Larson, E. (2024) “MNE-Python”. Zenodo. 10.5281/zenodo.14519545

25. Li, A., Feitelberg, J., Anand Prakash Saini, Höchenberger, R. and Mathieu Scheltienne (2022). MNE-ICALabel: Automatically annotating ICA components with ICLabel in Python. Journal of open source software, 7(76), pp.4484–4484. 10.21105/joss.04484.

26. Maister, L., Tang, T., & Tsakiris, M. (2017). Neurobehavioral evidence of interoceptive sensitivity in early infancy. eLife, 6. 10.7554/elife.25318

27. Obeid, I., & Picone, J. (2016). The Temple University Hospital EEG data corpus. Scholarshare. 10.34944/dspace/5136

28. Pang, J., Tang, X., Li, H., Hu, Q., Cui, H., Zhang, L., Li, W., Zhu, Z., Wang, J., & Li, C. (2019). Altered interoceptive processing in generalized anxiety disorder: A heartbeat-evoked potential research. Frontiers in Psychiatry, 10. 10.3389/fpsyt.2019.00616

29. Park, H.-D., & Blanke, O. (2019). Heartbeat-evoked cortical responses: Underlying mechanisms, functional roles, and methodological considerations. NeuroImage, 197, 502–511. 10.1016/j.neuroimage.2019.04.081

30. Plechawska-Wójcik, M., Augustynowicz, P., Kaczorowska, M., Zabielska-Mendyk, E., & Zapała, D. (2023). The influence assessment of artifact subspace reconstruction on the EEG signal characteristics. Applied Sciences, 13(3), 1605. 10.3390/app13031605

31. Pollatos, O., & Schandry, R. (2004). Accuracy of heartbeat perception is reflected in the amplitude of the heartbeat-evoked brain potential. Psychophysiology, 41(3), 476–482. 10.1111/1469-8986.2004.00170.x

32. Pollatos, O., Herbert, B. M., Mai, S., & Kammer, T. (2016). Changes in interoceptive processes following brain stimulation. Philosophical Transactions of the Royal Society B: Biological Sciences, 371(1708). 10.1098/rstb.2016.0016

33. Shamsaei, N., Shen, K. R., Wilder-Smith, E., Ong, C. J., & Li, X. (2010). Heartbeat evoked potential: A neural correlate of pain perception? Proceedings of the 32nd Annual International Conference of the IEEE Engineering in Medicine and Biology Society (EMBC), 1578–1581. 10.1007/978-3-642-14515-5_402

34. Schandry, R., Sparrer, B., & Weitkunat, R. (1986). From the heart to the brain: A study of heartbeat contingent scalp potentials. International Journal of Neuroscience, 30, 261–275. 10.3109/00207458608985677

35. Schmitz, M., Müller, L. E., Seitz, K. I., Schulz, A., Steinmann, S., Herpertz, S. C., & Bertsch, K. (2021). Heartbeat evoked potentials in patients with post-traumatic stress disorder: An unaltered neurobiological regulation system? European Journal of Psychotraumatology, 12(1). 10.1080/20008198.2021.1987686

36. Suksasilp, C., & Garfinkel, S. N. (2022). Towards a comprehensive assessment of interoception in a multi-dimensional framework. Biological Psychology, 168, 108262. 10.1016/j.biopsycho.2022.108262

37. Tahsili-Fahadan, P., & Geocadin, R. G. (2017). Heart–brain axis. Circulation Research, 120(3), 559–572. 10.1161/circresaha.116.308446

38. Terhaar, J., Viola, F. C., Bär, K.-J., & Debener, S. (2012). Heartbeat evoked potentials mirror altered body perception in depressed patients. Clinical Neurophysiology, 123(10), 1950–1957. 10.1016/j.clinph.2012.02.086

39. Ungureanu, M., Bigan, C., Strungaru, R., & Lazarescu, V. (2004). Independent component analysis applied in biomedical signal processing. Measurement Science Review, 4(2). 10.1088/0967-3334/26/1/r02

40. Yuan, H., Yan, H.-M., Xu, X.-G., Han, F., & Yan, Q. (2007). Effect of heartbeat perception on heartbeat evoked potential waves. Neuroscience Bulletin, 23(6), 357–362. 10.1007/s12264-007-0053-7

