## supplementary material for "Review of methods to derive the heartbeat-evoked potential: past practices and future directions"

Interaction between artefact settings and channels were investigated. Βeta (β) coefficients describe how the effect of different artefact correction settings on HEP values varies by channel, compared to the baseline, ICA without CFA removed setting. Results indicate the channels where a specific artefact setting significantly changes the HEP value (table 12). Most channels do not show significant interaction effects with artefact correction settings, suggesting that the artefact correction method does not strongly modulate HEP values across these channels. However, channels such as F3, F4, Fp1, and Fp2 exhibit significant differences in baseline (ICA without CFA removed) HEP values. Occipital and parietal EEG channels are significantly affected by artefact removal methods such as ASR-10 and ASR-5 and uncorrected EEG.

|  | Estimate | S.E. | Z val. | p |
| --- | --- | --- | --- | --- |
| ICA – F4 | -0.175 | 0.078 | -2.256 | 0.024 |
| ICA – F3 | 0.376 | 0.130 | 2.893 | 0.004 |
| ICA – F8 | -0.177 | 0.060 | -2.937 | 0.003 |
| ICA – Fp1 | 0.493 | 0.128 | 2.263 | 0.024 |
| ICA – Fp2 | -0.142 | 0.059 | -2.392 | 0.017 |
| ASR 10 – Cz | -0.097 | 0.045 | -2.146 | 0.032 |
| ASR 10 – Fp1 | -0.129 | 0.045 | -2.859 | 0.004 |
| ASR 10 – T3 | -0.097 | 0.045 | -2.145 | 0.032 |
| ASR 10 – T6 | -0.097 | 0.045 | -2.146 | 0.032 |
| ASR 5 – Fp1 | -0.343 | 0.045 | -7.549 | <0.001 |
| ASR 5 – O1 | -0.201 | 0.045 | -4.428 | <0.001 |
| ASR 5 – O2 | -0.210 | 0.045 | -4.625 | <0.001 |
| ASR 5 – P3 | -0.109 | 0.045 | -2.405 | 0.016 |
| ASR 5 – T3 | -0.190 | 0.045 | -4.189 | <0.001 |
| ASR 5 – T5 | -0.097 | 0.045 | -2.151 | 0.031 |
| ASR 5 – T6 | -0.135 | 0.045 | -2.964 | 0.003 |
| Uncorrected – C4 | 0.136 | 0.035 | 3.849 | <0.001 |
| Uncorrected – F3 | 0.180 | 0.035 | 5.098 | <0.001 |
| Uncorrected – Fp1 | 0.093 | 0.035 | 2.631 | 0.009 |
| Uncorrected – Fp2 | -0.421 | 0.035 | -11.929 | <0.001 |
| UncorrecteD – Fz | -0.521 | 0.035 | -14.764 | <0.001 |
| Uncorrected – O1 | 0.155 | 0.035 | 4.401 | <0.001 |
| Uncorrected – O2 | 0.077 | 0.035 | 2.173 | 0.030 |
| UncorrecteD – P3 | 0.183 | 0.035 | 5.196 | <0.001 |
| Uncorrected – P4 | 0.153 | 0.035 | 4.337 | <0.001 |
| Uncorrected – Pz | 0.104 | 0.035 | 2.950 | 0.003 |
| Uncorrected – T3 | 0.202 | 0.035 | 5.736 | <0.001 |
| Uncorrected – T5 | 0.083 | 0.035 | 2.347 | 0.019 |

**Table 1:** Significant interaction effects of artefact correction settings and channels on HEP values: linear mixed model results. S.E. – standard error; Z val. – Z value; P-unc - P uncorrected.

LMMs investigating the interaction between processing parameters and HEP were followed by post hoc analysis using estimated marginal means (EMMs) (table 2-7).

| Contrast | T | p-unc | p-corr | BF10 | hedges |
| --- | --- | --- | --- | --- | --- |
| 15 – 25 | -3.5851 | 0.0003 | 0.0223 | 12.337 | -0.0657 |
| 15 – 50 | -4.4216 | <0.001 | 0.0007 | 345.728 | -0.0808 |
| 20 – 24 | 4.0250 | <0.001 | 0.0038 | 63.508 | 0.0686 |
| 24 – 25 | -4.4022 | <0.001 | 0.0007 | 303.827 | -0.0744 |
| 24 – 30 | -3.4371 | 0.0006 | 0.0390 | 4.986 | -0.0513 |
| 24 – 45 | -3.6623 | 0.0003 | 0.0166 | 15.494 | -0.0620 |
| 24 – 50 | -5.1790 | <0.001 | <0.001 | 1.23E+04 | -0.0874 |
| 24 – 100 | -3.4610 | 0.0005 | 0.0356 | 7.512 | -0.0584 |
| 25 – 40 | 3.4735 | 0.0005 | 0.0340 | 7.928 | 0.05903 |
| 30 – 50 | -4.1876 | <0.001 | 0.0019 | 86.346 | -0.0392 |
| 35 – 50 | -3.7283 | 0.0002 | 0.0128 | 19.522 | -0.0625 |
| 40 – 50 | -4.3190 | <0.001 | 0.0010 | 210.354 | -0.0732 |

**Table 2:** EMM analysis for low-pass values used for HEP analysis.

| CONSTRAST | T | P-UNC | P-CORR | BF-10 | HEdges |
| --- | --- | --- | --- | --- | --- |
| 0.01 – 0.05 | -3.6435 | 0.0003 | 0.0057 | 15.509 | -0.0669 |
| 0.05 – 0.7 | 3.0916 | 0.0020 | 0.0419 | 2.3 | 0.0538 |
| 0.05 - 1 | 3.7458 | 0.0002 | 0.0038 | 21.598 | 0.0655 |

**Table 3:** EMM analysis for high-pass values used for HEP analysis.

| CONSTRAST | T | P-UNC | P-CORR | BF-10 | HEdges |
| --- | --- | --- | --- | --- | --- |
| 0.5 – 0.85 | 5.2011 | <0.001 | <0.001 | 1.36E+04 | 0.0866 |
| 0.5 – 0.88 | 7.9349 | <0.001 | <0.001 | 6.13E+11 | 0.0432 |
| 0.5 – 0.9 | 5.9323 | <0.001 | <0.001 | 7.93E+05 | 0.1005 |
| 0.55 – 0.85 | 5.1398 | <0.001 | <0.001 | 9935.66 | 0.0856 |
| 0.55 – 0.88 | 7.8076 | <0.001 | <0.001 | 2.25E+11 | 0.0425 |
| 0.55 – 0.9 | 5.8796 | <0.001 | <0.001 | 5.82E+05 | 0.0996 |
| 0.6 – 0.85 | 5.1657 | <0.001 | <0.001 | 1.14E+04 | 0.0860 |
| 0.6 – 0.88 | 7.8780 | <0.001 | <0.001 | 3.91E+11 | 0.0427 |
| 0.6 – 0.9 | 5.9018 | <0.001 | <0.001 | 6.63E+05 | 0.0999 |
| 0.65 – 0.85 | 5.0358 | <0.001 | <0.001 | 5862.645 | 0.0839 |
| 0.65 – 0.88 | 7.5342 | <0.001 | <0.001 | 2.77E+10 | 0.0414 |
| 0.65 – 0.9 | 5.7905 | <0.001 | <0.001 | 3.47E+05 | 0.0981 |
| 0.7 – 0.85 | 5.3440 | <0.001 | <0.001 | 2.89E+04 | 0.0890 |
| 0.7 – 0.88 | 7.7014 | <0.001 | <0.001 | 9.9E+10 | 0.0463 |
| 0.7 – 0.9 | 6.0574 | <0.001 | <0.001 | 1.68E+06 | 0.1025 |
| 0.75 – 0.85 | 4.7250 | <0.001 | 0.0001 | 1306.618 | 0.0789 |
| 0.75 – 0.88 | 5.4144 | <0.001 | <0.001 | 3.14E+04 | 0.0449 |
| 0.75 – 0.9 | 5.5147 | <0.001 | <0.001 | 7.41E+04 | 0.0930 |
| 0.8 – 0.85 | 3.5226 | 0.0004 | 0.0193 | 9.231 | 0.0587 |
| 0.8 – 0.88 | 3.7414 | 0.0009 | 0.0083 | 14.715 | 0.0262 |
| 0.8 – 0.9 | 4.4723 | <0.001 | 0.0004 | 410.377 | 0.0755 |

**Table 4:** EMM analysis for RR interval values used for HEP analysis.

| CONSTRAST | T | P-UNC | P-CORR | BF-10 | HEdges |
| --- | --- | --- | --- | --- | --- |
| 30 – 50 | -4.8090 | <0.001 | <0.001 | 2531.467 | -0.1010 |
| 50 – 70 | 4.2941 | <0.001 | 0.0008 | 199.188 | 0.07642 |
| 50 – 75 | 6.3367 | <0.001 | <0.001 | 9.95E+06 | 0.1147 |
| 50 – 100 | 7.2326 | <0.001 | <0.001 | 3.33E+09 | 0.0973 |
| 50 – 120 | 5.6979 | <0.001 | <0.001 | 2.16E+05 | 0.1000 |
| 50 – 150 | 4.7224 | <0.001 | 0.0001 | 1364.236 | 0.0840 |
| 75 – 80 | -4.4746 | <0.001 | 0.0003 | 428.946 | -0.0785 |
| 75 – 250 | -3.9968 | <0.001 | 0.0029 | 55.422 | -0.0675 |
| 80 – 100 | 4.9876 | <0.001 | <0.001 | 3569.665 | 0.0645 |
| 80 – 120 | 4.0495 | <0.001 | 0.0023 | 70.087 | 0.0698 |
| 100 – 250 | -4.6839 | <0.001 | 0.0001 | 778.898 | -0.0497 |
| 120 – 250 | -3.5401 | 0.0004 | 0.0181 | 9.903 | -0.0597 |

**Table 5:** EMM analysis for maximum epoch amplitude values used for HEP analysis.

| CONSTRAST | T | P-UNC | P-CORR | BF-10 | HEdges |
| --- | --- | --- | --- | --- | --- |
| -0.3 - -0.05 | -2.1532 | 0.0313 | 0.4698 | 0.193 | -0.0365 |
| -0.2 – -0.05 | -1.8907 | 0.0587 | 0.8806 | 0.081 | -0.0345 |
| -0.1 - -0.05 | -1.9136 | 0.0557 | 0.8353 | 0.119 | -0.0324 |

**Table 6:** EMM analysis for start of baseline correction window values used for HEP analysis.

| CONSTRAST | T | P-UNC | P-CORR | BF-10 | HEdges |
| --- | --- | --- | --- | --- | --- |
| 0.050 – 0.450 | -3.4792 | 0.0005 | 0.0227 | 8.047 | -0.0589 |
| 0.050 – 0.500 | -4.3996 | <0.001 | 0.0005 | 299.879 | -0.0745 |
| 0.100 – 0.400 | -3.3339 | 0.0009 | 0.0386 | 4.91 | -0.0565 |
| 0.100 – 0.450 | -3.9664 | <0.001 | 0.0033 | 49.183 | -0.0671 |
| 0.100 – 0.500 | -4.8554 | <0.001 | <0.001 | 2457.547 | -0.0822 |
| 0.150 – 0.400 | -3.5989 | 0.0003 | 0.01443 | 12.283 | -0.0609 |
| 0.150 – 0.450 | -4.2233 | <0.001 | 0.0011 | 140.463 | -0.0715 |
| 0.150 – 0.500 | -5.0951 | <0.001 | <0.001 | 8073.056 | -0.0863 |
| 0.200 – 0.450 | -3.8971 | <0.001 | 0.0044 | 26.969 | -0.0558 |
| 0.200 – 0.500 | -4.9888 | <0.001 | <0.001 | 3435.491 | -0.0747 |
| 0.250 – 0.450 | -3.5469 | 0.0004 | 0.0176 | 10.204 | -0.0601 |
| 0.250 – 0.500 | -4.4208 | <0.001 | 0.0004 | 329.128 | -0.0749 |
| 0.300 – 0.500 | -3.6631 | 0.0003 | 0.0113 | 15.501 | -0.0620 |

**Table 7:** EMM analysis for start of HEP window values used for HEP analysis.

| CONTRAST | T | P-UNC | P-CORR | BF-10 | HEdges |
| --- | --- | --- | --- | --- | --- |
| 0.250 - 0.600 | -4.0444 | <0.001 | 0.0024 | 48.398 | -0.0687 |
| 0.250 – 0.700 | -3.2630 | 0.0011 | 0.0497 | 3.888 | -0.0553 |
| 0.300 – 0.600 | -5.6374 | <0.001 | <0.001 | 1.08E+05 | -0.0847 |
| 0.300 – 0.650 | -3.6742 | 0.0002 | 0.0108 | 16.142 | -0.0622 |
| 0.300 – 0.700 | -4.3705 | <0.001 | 0.0006 | 263.962 | -0.0740 |
| 0.350 – 0.600 | -7.6617 | <0.001 | <0.001 | 7.47E+10 | -0.1094 |
| 0.350 – 0.650 | -5.0961 | <0.001 | <0.001 | 8114.769 | -0.0863 |
| 0.350 – 0.700 | -5.8455 | <0.001 | <0.001 | 4.83E+05 | -0.0990 |
| 0.400 – 0.600 | -6.6310 | <0.001 | <0.001 | 4.75E+07 | -0.0892 |
| 0.400 – 0.650 | -4.1545 | <0.001 | 0.0015 | 105.361 | -0.0704 |
| 0.400 – 0.700 | -4.9150 | <0.001 | <0.001 | 3285.371 | -0.0832 |
| 0.450 – 0.600 | -5.5653 | <0.001 | <0.001 | 7.18E+04 | -0.0747 |
| 0.450 – 0.650 | -3.3610 | 0.0008 | 0.0350 | 5.376 | -0.0569 |
| 0.450 – 0.700 | -4.1059 | <0.001 | 0.0018 | 86.225 | -0.0695 |
| 0.500 – 0.600 | -4.5394 | <0.001 | 0.0003 | 404.349 | -0.0591 |
| 0.500 – 0.700 | -3.2847 | 0.0010 | 0.0460 | 4.174 | -0.0556 |
| 0.550 – 0.600 | -3.9183 | <0.001 | 0.0040 | 29.299 | -0.0513 |

**Table 8:** EMM analysis for end of the HEP window values used for HEP analysis.
